# Guidance receptor-mediated mechanocompliance of GBM cells facilitates immune-silent invasion

**DOI:** 10.64898/2025.12.03.692092

**Authors:** Sangjo Kang, Chrystian Junqueira Alves, Haochen Tao, Christy Kolsteeg, Sita Sadia, Jack Y. Zhang, Jesse S. Fisher, Angela Dixon, Nadejda M. Tsankova, Aarthi Ramakrishnan, Li Shen, Roland H. Friedel, Hongyan Zou

## Abstract

The lethality of glioblastoma (GBM) stems from diffuse infiltration and immune evasion, two hallmarks traditionally studied separately. Here, we identify as unifying mechanism how GBM cells utilize guidance receptors Plexin-D1 and Plexin-B2 to gain mechanocompliance, i.e., the ability to deform and remodel membrane/cytoskeleton during confined migration without triggering immune activation. We show that *PLXND1* upregulation marks invasive fronts and predicts poor survival of glioma patients. Through live-cell imaging in microchannels, intracranial xenografts, single-nucleus transcriptomics, and lipidomics we demonstrate that Plexin-D1/B2 enable GBM cells to retract tumor microtubes (TMs), traverse constrictions, and escape microglial surveillance. Single and especially double deletion of *PLXND1* and *B2* resulted in TM overgrowth, membrane instability, and susceptibility to cell fragment shedding, leading to impaired migration and a shift to activation of tumor-associated myeloid cells. Our findings thus reveal a molecular strategy used by GBM cells to penetrate through interstitial space while escaping immune surveillance.

**SIGNIFICANCE:** Plexin-mediated mechanocompliance underlies the invasive yet immune-silent behavior of GBM cells, exposing a vulnerability that could be therapeutically exploited by forcing invading tumor cells into a mechanically fragile, immunogenic state.

## INTRODUCTION

Glioblastoma (GBM) remains uniformly fatal with limited treatment options ^1^. The high lethality stems from diffuse invasion, making complete surgical resection unattainable, and an immunologically “cold” tumor milieu refractory to new immunotherapies ^2^. So far, these hallmarks have largely been studied in isolation, yet invading GBM cells need to negotiate compact interstitial space while avoiding microglial detection. Understanding the missing link between confined migration and immune evasion will help uncover the molecular machinery underlying the invasive yet immune-silent migratory behavior of GBM cells, thus revealing new therapeutic targets to limit GBM spread while boosting immune activation.

GBM cells migrate preferentially along axonal tracts and microvessels ^3^. Recent studies revealed that GBM cells can extend long tumor microtubes (TMs) to form dense networks to synchronize activity and increase treatment resistance ^4^ and connect with neurons via pseudosynapses ^5^. Connected GBM cells at tumor core migrate significantly slower than unconnected cells and the connected state is inversely associated with tumor invasion ^6^. Thus, initiation of invasion and switching from stationary to infiltrative state may require GBM cells to retract TMs and enhance membrane turnover to remodel adhesions to neighboring cells and extracellular matrix (ECM). Moreover, mechanical stress during confined migration risks membrane tearing and cell shearing, potentially flagging cells to microglia, resident immune cells that constantly survey the brain microenvironment. Hence, invasive GBM cells need to acquire mechanoplasticity, i.e., the ability to deform, repair, and remodel membrane/cytoskeleton under confinement for safe passage through constrictions while reducing immune visibility.

Plexins are conserved transmembrane receptors originally identified as axon guidance molecules during neurodevelopment ^7^. Mammalian genomes contain nine plexins divided into four classes: Plexins A1-A4, B1-B3, C, and D1 ^8^. The ring-shaped extracellular domain of plexins binds to secreted or membrane-bound semaphorins ^9^, leading to activation of the conserved intracellular GAP (GTPase activating protein) domain, which in turn deactivates Rap/R-Ras small GTPases to finetune cytoskeleton dynamics and cell adhesion ^10–13^. Beyond their roles in neurodevelopment, plexins also regulate a wide range of cellular processes in adult physiology of the central nervous, immune, and vascular systems ^14–17^. Aberrant expression of plexins and semaphorins are linked to increased malignancy of various cancers ^18–20^. We recently showed that Plexin-B2 promotes GBM invasiveness by reducing cell adhesion, enhancing actomyosin contractility, and orchestrating migratory behavior in response to matrix stiffness and confined space ^19, 21, 22^. Plexin-D1, best known in vascular biology and recently implicated as a mechanosensor responding to shear force of blood flow in endothelial cells ^23^, is upregulated in many cancers, in both tumor cells and tumor vasculature ^24^. These findings prompted us to ask whether Plexin-D1 cooperates with Plexin-B2 to endow GBM cells with mechanocompliance necessary for immune-silent GBM invasion.

Here, we show that Plexin-D1/B2 are required for GBM cells to retract long processes, disengage from TM networks, and passage through constrictions without physical damage. Deletion of plexins turns tumor cells into a mechanically fragile and immunogenic state, not only impairing migration but also provoking microglial activation. Understanding the molecular strategies used by invading GBM cells to maintain mechanocompliance for immune-silent invasion points to new strategies to curb GBM invasion while increasing immune activation.

## RESULTS

### Plexin-D1 is enriched at GBM borders and associates with poor patient survival

To identify molecular drivers of GBM invasion, we surveyed the TCGA glioma patient database for Plexin expression, which revealed higher levels of *PLXND1* and *PLXNB1-3* as compared to *PLXNA1-4* and *PLXNC1* (**Fig. 1a**). Analysis of both TCGA and CGGA glioma databases also revealed that *PLXND1* is expressed at higher levels in high-grade versus low-grade gliomas and associates with poor survival, similar to *PLXNB2* ^21^ (**Fig. 1b**; **Fig. S1a-c**).

**Figure 1.**
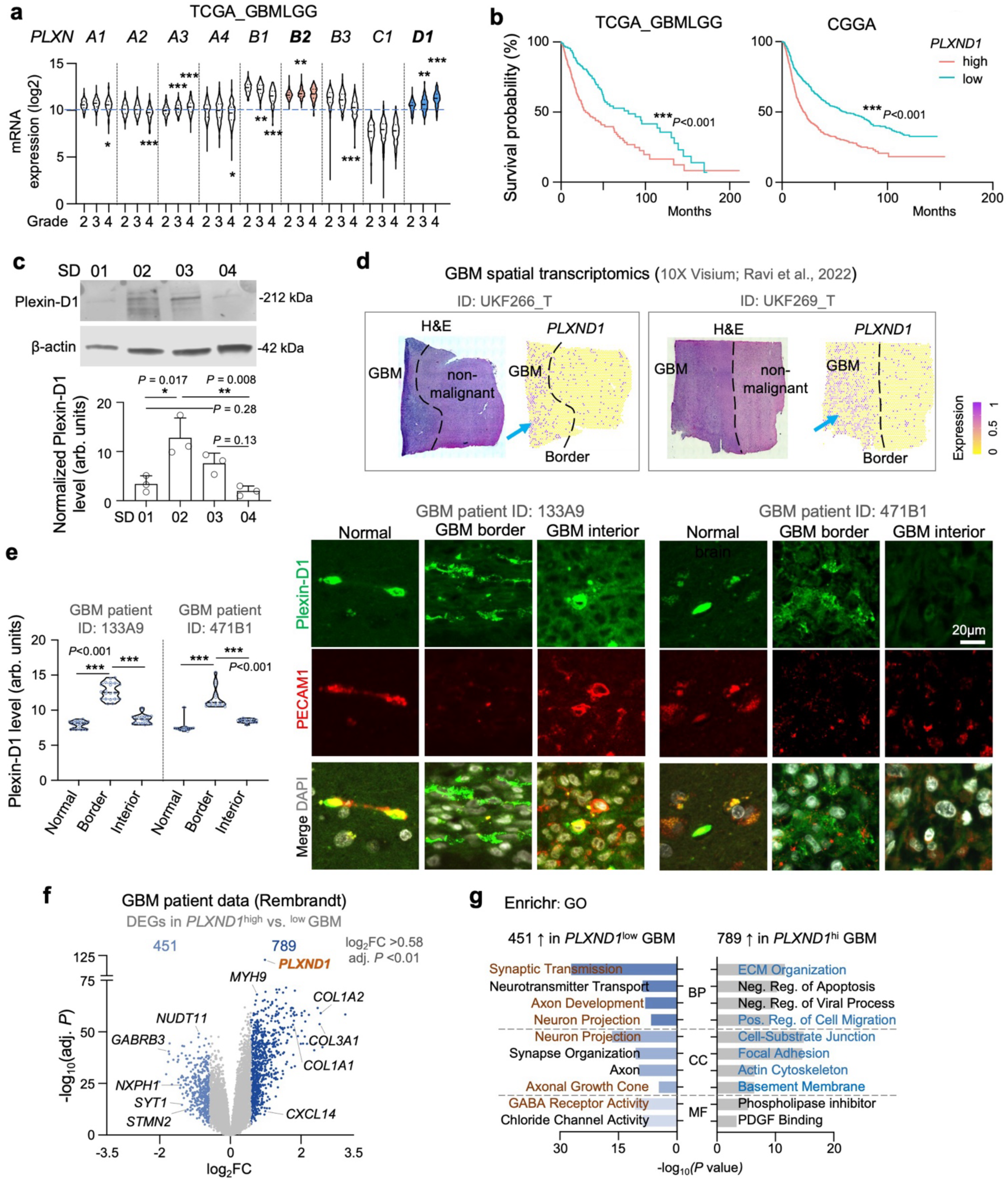
Plexin-D1 is upregulated in high grade gliomas and associated with poor outcome. **a)** Violin plots show high expression of *PLXNB2* and *PLXND1* in gliomas, with increasing levels in higher grades (TCGA_GBMLGG dataset at GlioVis platform). Median and quartiles are shown inside violins. **b)** High expression of *PLXND1* correlates with poor patient poor survival in both TCGA and CGGA human glioma data sets. Patients were stratified by median expression levels of *PLXND1*. **c)** Western blot for Plexin-D1 protein in four patient-derived GSC lines, with β-actin as loading control. For quantification, n=3 independent experiments. Bar graphs represent mean ± s.e.m. Two-way ANOVA with Tukey’s multiple comparisons test. **d)** Spatial transcriptomics data from two GBM patient samples show *PLXND1* expression at GBM border. **e)** IF analysis of two human GBM surgical specimens reveals higher Plexin-D1 expression at GBM borders compared to interior. Note that in normal brain tissue, Plexin-D1 co-localizes with vascular marker PECAM1. n=15-20 imaged regions for each group. Violin plots indicate median and quartiles. One-way ANOVA with Dunnett’s multiple-comparison test. **f)** Volcano plot shows differentially expressed genes (DEGs) of human GBMs stratified by high vs. low expression of *PLXND1* (split by median; Rembrandt GBM dataset at GlioVis). **g)** Enriched gene ontologies (GO) of up- or down-regulated DEGs in human GBMs expressing high vs. low *PLXND1*. BP, biological processes; CC, cellular component; MF, molecular function.

Western blotting confirmed Plexin-D1 protein expression in all four patient-derived IDH1 wildtype glioma stem cell (GSC) lines that our laboratory has previously characterized ^19^ (**Fig. 1c**). Consistent with the intertumoral heterogeneity of GBM, SD2 (mesenchymal subtype) and SD3 (proneural) GSCs expressed higher levels of Plexin-D1 than SD1 (classical) and SD4 (proneural) GSCs.

To resolve spatial patterns of Plexin-D1 expression, we surveyed spatial transcriptomics data of GBM patient samples ^25^, which revealed an enrichment of *PLXND1* expression at tumor borders (**Fig. 1d**). For validation, we conducted immunofluorescence (IF) staining of GBM patient specimens from the Mount Sinai brain bank, which confirmed elevated Plexin-D1 expression at GBM borders, in contrast to tumor interior and normal brain areas where Plexin-D1 was mainly detected on vascular cells (**Fig. 1e**).

Analysis of differentially expressed genes (DEGs; |log_2_FC|>0.58, adj. *P*<0.01) in human GBMs stratified into *PLXND1*^high^ vs. *PLXND1*^low^ groups revealed upregulation of genes associated with ECM organization, cell migration, and cell-substrate junction, and downregulation of neuronal projection/synaptic gene modules (**Fig. 1f, g**). Collagen genes, such as *COL1A1*, a marker of collectively migrating GBM cells ^26^, and *COL1A2*/*3A1*, were among the genes most correlated with *PLXND1* expression (**Fig. 1f**; **Fig. S1d**).

Together, these GBM patient data on Plexin-D1 upregulation at invasive fronts, correlation with poor survival, and co-upregulation of genes involved in cell migration and ECM organization and co-downregulation of genes concerning cell projections support a role of Plexin-D1 as driver of GBM invasion.

### Plexin-D1 deletion impairs confined migration and reduces membrane turnover

To explore the function of Plexin-D1 in GBM cells, we generated next GSC lines with CRISPR/Cas9 knockout of *PLXND1* (*PD1* KO) (**Fig. S2a**), using Cas9 with sgRNA against *GFP* for control studies. The loss of Plexin-D1 protein in GSCs was verified by Western blotting and IF (**Fig. 2a, b**).

**Figure 2.**
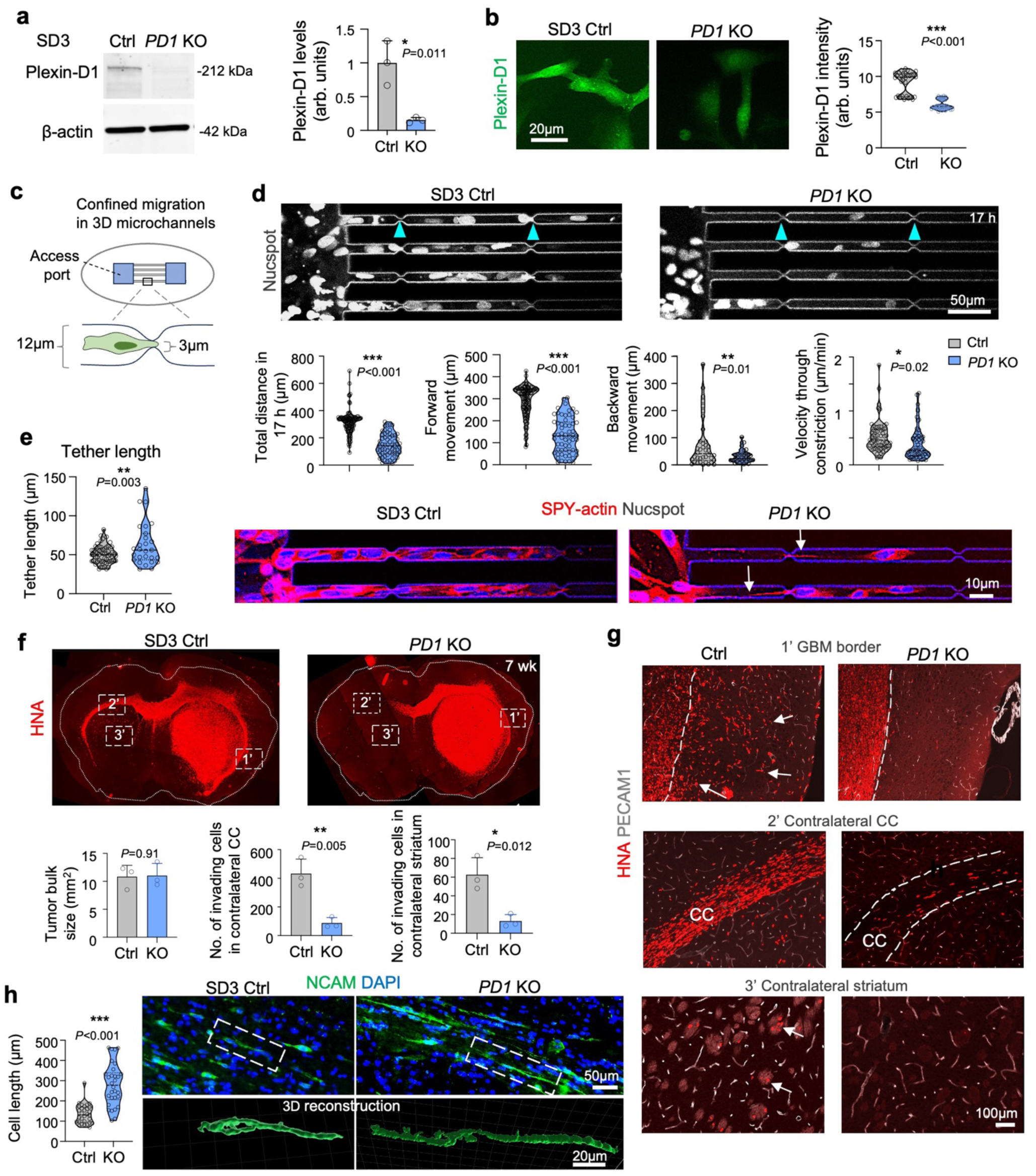
Plexin-D1 ablation impedes GBM invasion. **a)** Western blot shows reduction of Plexin-D1 protein in SD3 GSCs with CRIPSR knockout of *PLXND1* (*PD1* KO). SD3 cells carrying CRISPR/Cas9 vector with sgRNA against GFP served as negative control. β-actin as loading control. n=3 independent experiments. Bar graphs represent mean ±s.e.m. Unpaired two-tailed Student’s t-test. **b)** SD3 *PD1* KO GSCs show reduced Plexin-D1 protein expression by IF compared to control cells. n=30 cells for each condition. Violin plots showing median and quartiles. Unpaired two-tailed Student’s t-test. **c)** Diagram of confined migration assay in 12 µm-wide microchannels with periodic 3 µm constrictions. **d)** Live cell imaging of GSCs labeled the nuclear dye Nucspot at 17 h after seeding. Note drastic reduction of *PD1* KO cells that traverse the microchannels with constrictions (arrowheads). n>40 cells for each group, from 3 independent experiments. Violin plots show median and quartiles. Unpaired two-tailed Student’s t-test. **e)** Migrating *PD1* KO cells in microchannels formed long tethers between cells (arrows). n=27 and 60 cells for Ctrl and *PD1* KO, respectively; from 3 independent experiments. Violin plots showing median and quartiles. Unpaired two-tailed Student’s t-test. **f, g)** Coronal sections (f) of mouse brains stained for human nuclear antigen (HNA) at 7 wk post-transplant show comparable bulk size of *PD1* KO and control SD3 tumors but reduced diffuse infiltration at GBM border (dashed line) and less invasion along corpus callosum (CC) and into contralateral striatum. Enlarged images of boxed areas are shown in (g). n=3 mice per group. Bar graphs represent mean ± s.e.m. Unpaired two-tailed Student’s t-test. **h)** High magnification images and 3D reconstructions (Imaris) reveal elongated morphology *PD1* KO GBM cells (stained by IF for human NCAM) extending long processes during migration along corpus callosum. n=30 cells for each condition, from three independent transplants. Violin plots show median and quartiles. Unpaired two-tailed Student’s t-test.

Plexin-D1 KO deletion did not affect proliferation or 2D scratch closure (**Fig. S2b-d**). However, in 3D microchannel devices with laminin-coated glass floors and silicon walls (10 µm height, 12 µm width, periodic 3 µm constrictions), time-lapse imaging of *PD1* KO cells exhibited reduced forward and backward movements, as well as lower velocity through constrictions (**Fig. 2c, d**; **Video S1**). Of note, the migration of GSCs into the microchannels occurred spontaneously, as no chemoattractants were added. Interestingly, we also observed formation of F-actin containing thin long tethers linking neighboring *PD1* KO cells, suggesting incomplete detachment of stretched cell processes (**Fig. 2e**; **Video S2**). The impaired confined migration and the thin long tethers are also reminiscent of Plexin-B2 KO phenotypes in GSCs ^22^.

We further studied the effect of Plexin-D1 deletion on the dynamics of the plasma membrane by live-cell imaging with fluorescent probes. First, we found that the membrane-anchored reporter myrPalm-CFP was mainly localized on intracellular vesicles/endosomes in control GSCs at 3 days after transfection but was found predominantly retained on the plasma membrane in *PD1* KO cells, suggesting reduced endocytosis in mutant cells (**Fig. S2e**). Second, imaging of fluorescent reporter PH-PLCδ1-GFP that binds to phosphatidylinositol 4,5-bisphosphate (PIP2) showed similar results (**Fig. S2f**). Third, signal intensity of FluoVolt, a voltage-sensitive membrane dye, revealed a drastic decrease in *PD1* KO cells, signifying a more negative inner membrane charge (**Fig. S2g**). These phenotypes again mirror the membrane features of Plexin-B2 KO GSCs ^22^.

When examining the morphology of cultured GSCs, we found that *PD1* KO cells frequently extended thin long processes containing F-actin, tubulin and integrin β1; mitochondria also populated these projections, but lysosome content was low (**Fig. S3a, b**). These thin long processes touched neighboring cells, resembling the long tethers seen in 3D microchannels and the tumor microtubes (TM) that connect GBM cells in vivo ^27^.

### Plexin-D1 loss limits intracranial spread and shifts migratory routes of GBM cells

To examine the functional impact of Plexin-D1 on GBM progression in vivo, we transplanted SD3 GSCs into the striatum of SCID mice. Both control and *PD1* KO cells formed similar tumor bulk at 7 weeks post-transplant, but *PD1* KO tumors exhibited less diffuse invasive margins and reduced spreading into the contralateral hemisphere along the corpus callosum (**Fig. 2f, g**). Reminiscent of the long tethers seen in 3D microchannels and the TM-like processes in cultures, invading *PD1* KO cells in corpus callosum displayed an elongated morphology revealed by IF for human NCAM to outline cell contours and 3D rendering of cell surfaces (**Fig. 2h**). Notably, loss of Plexin-D1 also shifted migratory paths from striatal axon bundles toward perivascular routes (**Fig. S4a, b**), similar to Plexin-B2 KO phenotypes in vivo ^19^.

When examining the association of myeloid cells (PU.1^+^) with invading tumor cells, we observed a significantly higher ratio of PU.1^+^ cells relative to tumor cells at invasion fronts and along invasion paths in the corpus callosum in *PD1* KO tumors (**Fig. S5a, b**). This observation suggests that the reduced mechanocompliance by *PD1* KO (TM-like processes, long tethers, stretched out morphology) may increase immune visibility of invading cells.

### Single-nucleus RNA-seq shows depletion of invasives state in *PD1* KO GBM

We next conducted single-nucleus (sn) RNA-seq on SD3 GBM xenografts at 7 week post intracranial transplant, yielding after quality control data from 19,417 nuclei (n = 2 per genotype; **Fig. 3a**; **Fig. S6a, b**). Human GBM cells were readily distinguishable from mouse stromal cell populations by marker gene expression (**Fig. S6c**).

**Figure 3.**
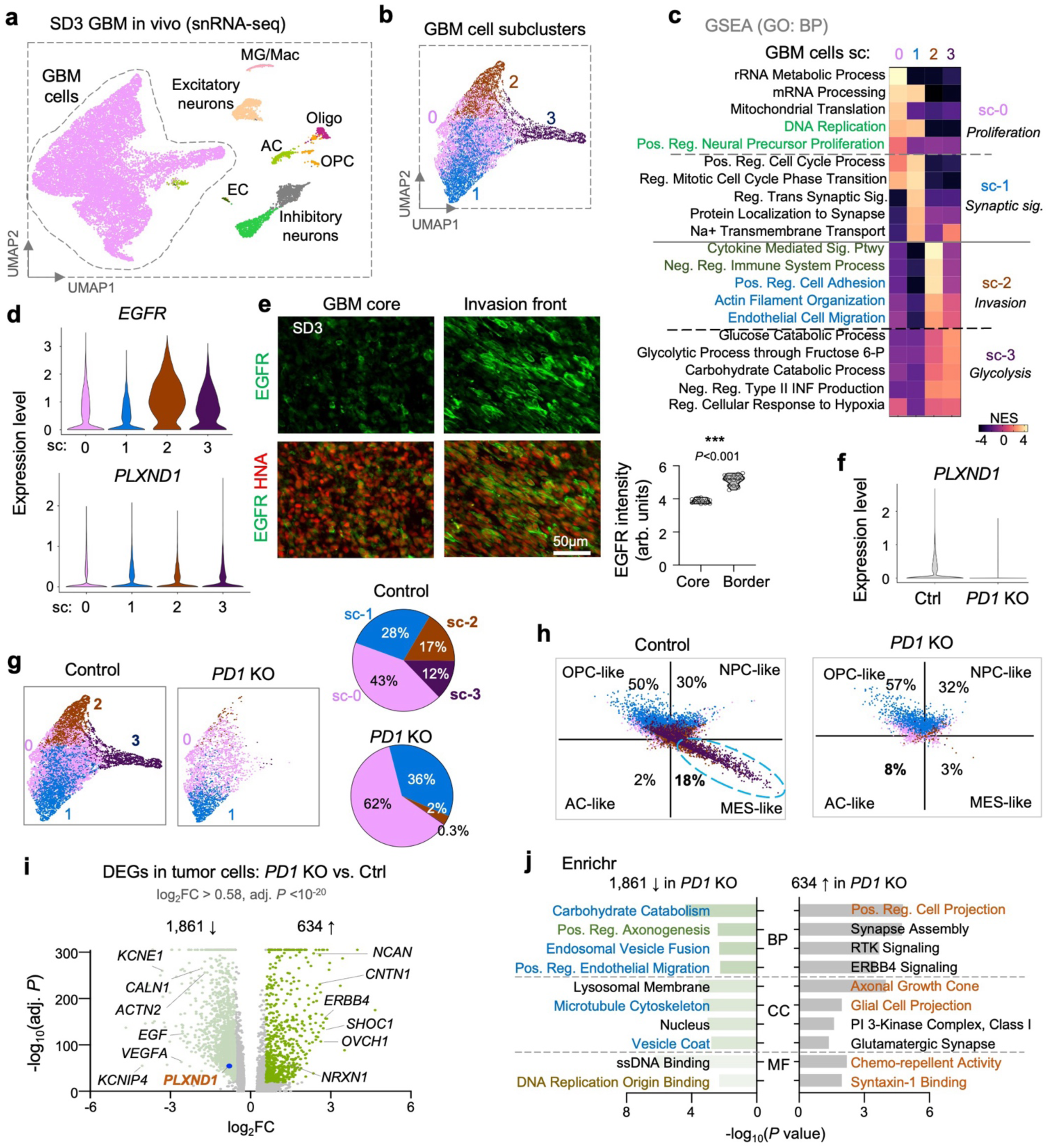
Single cell transcriptomics highlights reduction of invasive GBM subclusters. **a)** UMAP embedding of snRNA-seq data from control and *PD1* KO SD3 tumors at 7-week post-transplant (from n=2 samples per condition), with major cell types annotated. **b)** UMAP embedding of the GBM tumor cells only, with four subclusters indicated. **c)** Enriched gene ontologies for Biological Process (GO: BP) across GBM cell subclusters (sc). Note that sc0/sc1 feature proliferation and synaptic signaling, while sc2/sc3 feature invasion and glycolysis. **d)** Violin plots show *EGFR* and *PLXND1* expression in four GBM cell subclusters. **e)** IF of brain tumor sections reveals higher EGFR expression at GBM invasion front than in GBM core of SD3 xenograft tumors. Quantifications from n=15 regions from three independent transplants. Violin plots showing median and quartiles. Unpaired two-tailed Student’s t-test. **f)** Reduction of *PLXND1* expression in SD3 *PD1* KO tumor cells compared to control xenografts. **g)** UMAP embedding of GBM tumor cells, separated by genotypes. Note the drastic depletion of sc2 and sc3 in *PD1* KO GBM, visualized by pie chart. **h)** Analysis of Neftel et al. GBM cell states shows that mesenchymal (MES)-like state was reduced from 18% to 3%, whereas astrocyte (AC)-like was expanded from 2% to 8% in *PD1* KO tumor compared to control tumor. **i)** Volcano plot shows DEGs of GBM cells of *PD1* KO vs. control tumors based on snRNA-seq dataset. **j)** Enrichr gene ontology analysis of DEGs from GBM cells of *PD1* KO vs. control tumors. BP, biological processes; CC, cellular component; MF, molecular function.

GBM cells were resolved into four subclusters (sc), with sc-0 characterized by gene signatures associated with a proliferative state, sc-1 with synaptic organization, and sc-2/3 with invasive/glycolytic states (**Fig. 3b, c**; **Fig. S6d-f**). Consistently, sc-2/3 expressed higher levels of *EGFR,* a growth factor receptor frequently upregulated in malignant gliomas ^28^ (**Fig. 3d**). IF confirmed higher EGFR expression at invasion fronts of SD3 GBM, supporting a link of sc-2/3 with invasive state (**Fig. 3e**).

*PLXND1* was expressed at comparable levels across four tumor cell subclusters with a trend of increase in sc-2/3 (**Fig. 3d**). We confirmed reduced *PLXND1* expression in *PD1* KO tumors (**Fig. 3f**). Remarkably, *PD1* KO tumors were depleted for invasive cell states, with sc-2/3 shrinking from ∼17%/12% in control tumors to ∼3%/0.3% in *PD1* KO tumors (**Fig. 3g**). Scoring for GBM cell states according to signatures defined by Neftel and colleagues ^29^ showed a clear drop in mesenchymal-like cells (18%→3%) in *PD1* KO tumors (**Fig. 3h**), signifying a shift away from invasive states.

Echoing the enriched gene ontologies (GOs) of the DEGs of *PLXND1*^high^ vs. *PLXND1*^low^ GBM patients (see Fig. 1), upregulated DEGs in GBM cells from *PD1* KO vs. control xenografts featured GO terms associated with cell projection and axon growth-cone, while downregulated DEGs featured microtubule cytoskeleton, endosomal vesicle fusion, and carbohydrate catabolism, consistent with suppressed membrane/cytoskeletal remodeling and invasive state, but enhanced TM-like projections and connected state in KO (**Fig. 3i, j**).

Hence, snRNA-seq data converge with the KO phenotypes observed by live-cell imaging, microchannel assays, and in vivo intracranial xenografts, supporting the model that Plexin-D1 upregulation switches GBM cells from a connected/stationary state to an unconnected/invasive state.

### Double Plexin-D1/B2 deletion drives TM outgrowth, mechanical fragility and impaired confined migration

Since Plexin-D1 KO phenocopies Plexin-B2 KO in many aspects, including impaired confined migration, reduced GBM invasiveness, shift of migratory path, and TM-like long processes ^19, 22^, we next probed functional redundancy by generating CRISPR/Cas9 double knockouts (*PB2*/*PD1* DKO) in both SD2 and SD3 GSC lines (**Fig. 4a**). IF confirmed loss of Plexin-B2 and -D1 proteins in DKO cells (**Fig. 4b**).

**Figure 4.**
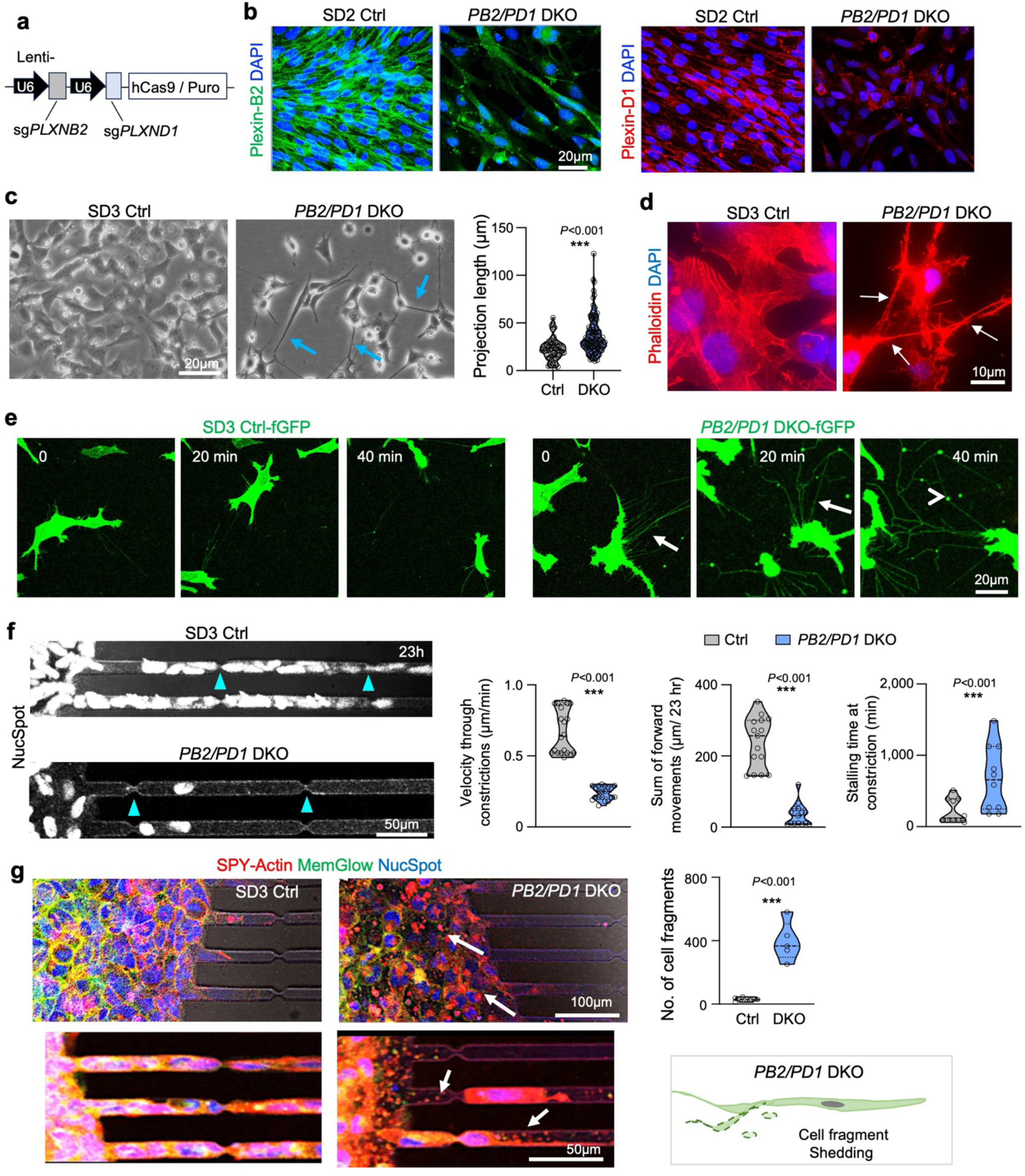
Double deletion of Plexin-D1 and -B2 limits withdrawal of processes and impairs confined migration. **a)** Diagram of lentiviral vector with two sgRNAs targeting both *PLXND1* and *PLXNB2* for double knockout (*PB2/PD1* DKO). **b)** Reduced Plexin-D1 and -B2 protein expression by IF in cultured SD2 *PB2/PD1* DKO GBM cells compared to controls. **c)** Phase contrast images show long projections from SD3 *PB2/PD1* DKO cells that resemble tumor microtubes (arrows). n=60-120 cells from three independent cultures for each condition. Violin plots show median and quartiles. Unpaired two-tailed Student’s t-test. **d)** Phalloidin staining shows presence of F-actin inside the tumor microtubes (arrows). **e)** Time-lapse live-cell imaging of SD3 GBM cells expressing membrane anchored fGFP. Note that *PB2/PD1* DKO cells display long extended tumor microtubes (arrows) and cell fragment shedding (arrowhead). **f)** Confined migration assay shows that SD3 *PB2/PD1* DKO GSCs have severely hampered confined migration capacity. Arrowheads point to constrictions. n=20 cells for velocity, n=15 cells for forward movement n=10 for stalling time at constriction from two independent assays for each condition. Violin plots show median and quartiles. Unpaired two-tailed Student’s t-test. **g)** Live-cell imaging of migrating GBM cells in microchannels cells labeled with SPY-actin, MemGlow and NucSpot. Note numerous cell fragments of *PB2/PD1* DKO GSCs inside of access port and microchannels (arrows). n=5 experiments for each condition. Unpaired two-tailed Student’s t-test. Violin plots show median and quartiles. Schematic on right.

When examining cell morphology of GSCs cultured on laminin, we observed that DKO cells formed abundant, long TM networks rich in F-actin and pMLC2 (**Fig. 4c, d**; **Fig. S7a, b**), while cell proliferation was also negatively affected (**Fig. S7b, c**). DKO cells also showed increased expression of GAP-43 (**Fig. S7b)**, which is involved in TM formation ^27^. FluoVolt signals were significantly decreased, indicating altered local membrane electric charge (**Fig. S7e**). Live-cell imaging with NR12A membrane dye revealed increased membrane ruffling in DKO cells, likely linked to reduced overall membrane tension (**Fig. S7f; Video S3**). Live-cell imaging with the membrane anchored farnesylated (f)GFP reporter further revealed numerous thin long processes projecting from DKO cells that formed a TM network (**Fig. 4e**). Videography also revealed frequent TM retractions and ruptures, generating fGFP^+^ cellular debris, indicating mechanical fragility (**Fig. 4e**). Imaging with mitotracker and SPY-tubulin probes revealed the presence of mitochondria and tubulin in the TM-like projections; we also observed withdrawal of TMs filled with mitochondria, but no evidence of mitochondria exchange between neighboring cells through TMs during a 1 hr timeframe (**Fig. S7g**; **Videos S4-S5**).

In microchannel assays, invading *PB2*/*PD1* DKO cells showed markedly slower velocity through constrictions, longer stalling times, and reduced sum of forward movements (**Fig. 4f**). In addition, in the entrance port and inside the channels, we observed abundant MemGlow/SPY-actin^+^ fragments from *PB2/PD1* DKO cells (**Fig. 4g**; **Video S6**), indicating shedding.

### Double Plexin-D1/B2 deletion affects membrane-related transcriptomes and lipidomes of GSCs

To understand the transcriptional consequences of single or double deletion of Plexin-D1/B2, we conducted RNA-seq of GSCs (**Fig. S8a**). For both SD2 and SD3 GSCs, RNA-seq samples were clearly separated by genotypes after principal component analysis (PCA), with PC1 capturing ∼83% of variance for SD3 DKO and ∼78% for SD2 DKO, while PC2 captured ∼16% and ∼21% for single KO in SD3 and SD2, respectively (**Fig. 5a, d**). This indicates transcriptional adaptations to Plexin-D1/B2 loss, with a more profound change upon *PB2/PD1* DKO.

**Figure 5.**
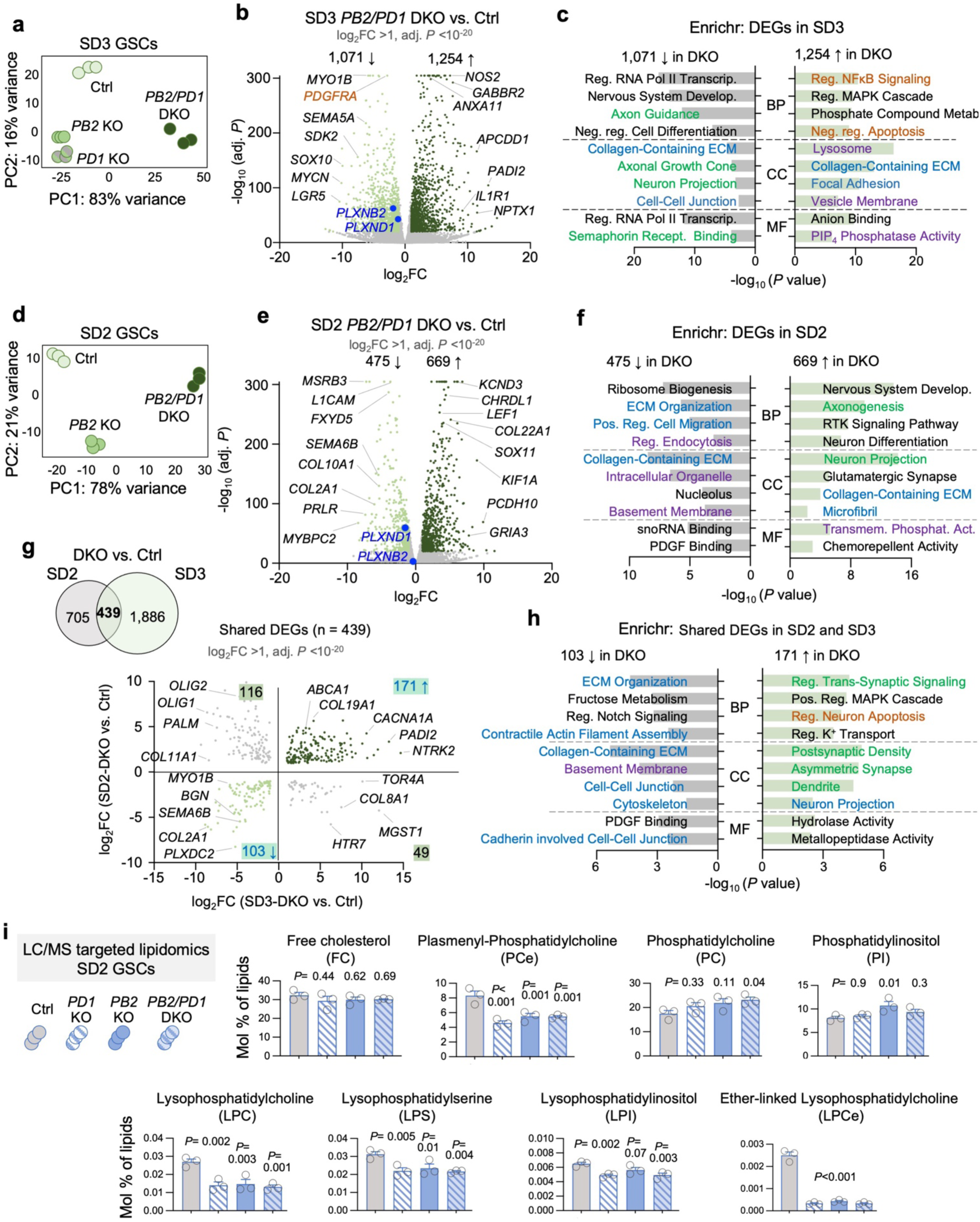
Double deletion of Plexin-D1/B2 alters transcriptomic and lipid profiles of GSCs related to membrane dynamics. **a)** Principal component (PC) analysis of RNA-seq data of three independent samples of cultured SD3 GSCs for each genotype shows that *PB2/PD1* DKO samples segregated from others along PC1, and single KOs segregated from controls along PC2. **b)** Volcano plot shows DEGs of *PB2/PD1* DKO cells vs. controls in SD3. **c)** Enrichr analysis of DEGs of *PB2/PD1* DKO vs. control SD3 cells. BP, biological processes; CC, cellular component; MF, molecular function. **d)** PCA of RNA-seq of SD2 GSC line shows segregation of *PB2/PD1* DKO samples along PC1, and of single *PB2* KO along PC2. **e)** Volcano plot shows DEGs of SD2 *PB2/PD1* DKO cells vs. control. **f)** Enrichr analysis of the DEGs of SD2 *PB2/PD1* DKO vs. control GSCs. **g)** Top, Venn diagram shows overlap of *PB2/PD1* DKO DEGs of SD2 and SD3 GSCs. Bottom, shared DEGs were plotted into quadrants according to changes in SD2 and SD3 cells. **h)** Enrichr analysis of the shared DKO DEGs with concordant changes in SD2 and SD3 *PB2/PD1* DKO cells. **i)** Targeted lipidomics analysis reveals changes in relative lipid content of SD2 mutant cells with single or double deletion of Plexin-D1/B2 compared to control cells. For quantification, n=3 independent cultures. Bar graphs represent mean ± s.e.m. One-way ANOVA with Dunnett’s multiple-comparison test.

In SD3, using stringent cutoffs (|log_2_FC| > 1 and adj. *P* <10^-20^), we identified 1,254 upregulated and 1,071 downregulated DEGs in *PB2/PD1* DKO vs. control cells. DKO cells upregulated genes for focal adhesion/ECM/vesicle-membrane/lysosome and PIP-phosphatase, and downregulated genes for axon guidance/semaphorin signaling (confirming genotype) and cell-cell junctions (**Fig. 5b, c**). We also compared the DEGs of *PD1* or *PB2* single KO vs. control SD3, which interestingly showed significant overlap, with 650 shared DEGs with concordant directionality (**Fig. S8b-g**). These DEGs regulate cell migration, vesicle membrane, cell projection, and ECM organization.

Similarly, in SD2, *PB2/PD1* DKO upregulated DEGs (n=669) for axonogenesis/neuron projection, collagen/ECM, microfibril, and transmembrane phosphatase activity, and downregulated DEGs (n=475) for cell migration and membrane processes (regulation of endocytosis, intracellular organelle) (**Fig. 5e, f**).

Intersection analysis revealed that 439 DEGs in DKO were shared by SD3 and SD2, with majority (62%) showing concordant directionality (**Fig. 5g**). The shared upregulated DEGs were enriched for synaptic signaling and neuron projection, and the shared downregulated DEGs featured contractile actin assembly (**Fig. 5h**).

Next, we performed targeted LC-MS lipidomics in SD2 GSCs, with triplicate samples for control, single or double KO of Plexin-D1/B2 (**Fig. 5i**). This identified a wide range of changes of lipid components in both single and double KO relative to controls. While free cholesterol (FC) levels were not significantly changed, indicating no major changes in sterol metabolism, we observed a decrease of plasmenyl-phosphatidylcholine (PCe) and a slight increase of phosphatidylcholine (PC) as well as phosphatidylinositol (PI) species, consistent with a more negatively charged inner leaflet (**Fig. 5i**). Interestingly KO and double KO cells also showed reduced levels of several species of lysophopholipids, including lysophophatidycholine (LPC), lysophosphatylserine (LPS), lysophosphatidylinosotol (LPI) and ether-linked lysophosphotidylcholine (LPCe), suggesting a decrease in phosopholipase A2 activation and less local membrane remodeling (**Fig. 5i**). Together, these lipidomics findings are in line with altered endocytosis vesicle/membrane turnover and a less mechanocompliant plasma membrane upon Plexin ablation.

Altogether, transcriptomics and lipidomics data add molecular evidence to the model that Plexin-D1 and -B2 regulate membrane turnover and cytoskeletal remodeling that allows GBM cells to withdraw long processes, initiate detachment from TM networks, and migrate through confined space without sustaining physical damage or shedding of cell fragments.

### Double deletion of Plexin-D1/B2 limits GBM spread and heightens immune activation

To study the in vivo consequences of the altered membrane properties of *PB2/PD1* DKO cells, we conducted intracranial transplant studies. *PB2/PD1* DKO tumors were markedly smaller than controls and tumor spread was drastically reduced (**Fig. 6a, b**; **Fig. S9a**). Invading *PB2/PD1* DKO cells in the corpus callosum were extensively elongated with stretched long processes (**Fig. 6c**). PDGF receptor α expression was diminished in both tumor core and edge of *PB2/PD1* DKO tumors (**Fig. 6d**), consistent with RNA-seq data identifying *PDGFRA* as one the most significant downregulated DEGs in *PB2/PD1* DKO (see Fig. 5a).

**Figure 6.**
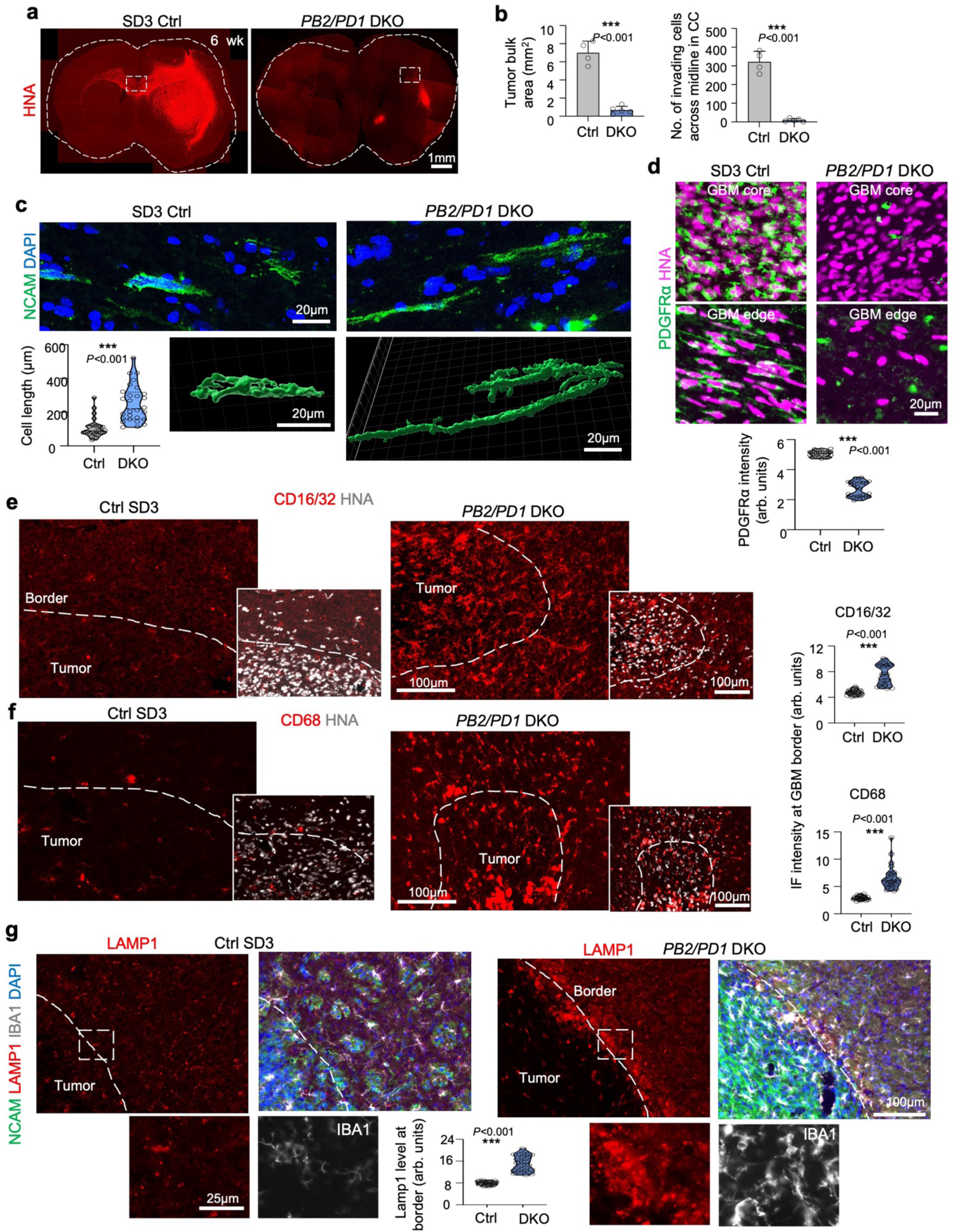
Reduced invasion and enhanced immune activation in GBM with Plexin-B2/D1 double deletion. **a)** Overview of SD3 control and *PB2/PD1* DKO tumor expansion, visualized by IF for human nuclear antigen (HNA) at 6 weeks post-transplant. **b)** Quantification of tumor bulk area and the number of invading cells across midline show reduction in *PB2/PD1* DKO transplants. n=4 for control and n=5 for *PB2/PD1* DKO, bar graphs represent mean ± s.e.m. Unpaired two-tailed Student’s t-test. **c)** IF images show elongated shape of SD3 *PB2/PD1* DKO cells during migration in corpus callosum. 3D renderings (bottom) highlight long processes of DKO cells. n=28 cells from 3 independent transplants for each group. Violin plots show median and quartiles. Unpaired two-tailed Student’s t-test. **d)** IF images show that in SD3 *PB2/PD1* DKO tumors, tumor cells at both core and edge express less PDGF receptor α. n=27 regions from 3 independent transplants for each group. Violin plots show median and quartiles. Unpaired two-tailed Student’s t-test. **e, f)** IF images show that CD16/32 and CD68 were elevated in SD3 *PB2/PD1* DKO tumors. n=30 cells from 3 independent transplant for each group. Violin plots show median and quartiles. Unpaired two-tailed Student’s t-test. **g)** IF images at tumor borders in striatum show enhanced LAMP1 expression in SD3 *PB2/PD1* DKO tumors. Enlarged images of boxed area are shown at bottom. n=30 regions from 3 independent transplants for each group. Violin plots show median and quartiles. Unpaired two-tailed Student’s t-test.

Notably, myeloid cells appeared markedly activated in *PB2/PD1* DKO tumors at both GBM borders and invasion paths, evidenced by higher IBA1 intensity and larger size of IBA^+^ cells surrounding invading tumor cells (**Fig. S9b**). Co-IF for PU.1 and human NCAM also revealed increased cell ratios of PU.1^+^ myeloid cells to invading GBM cells in the corpus callosum (**Fig. S9c**). DKO tumors also displayed increased expression of CD16/CD32 and CD68, both inflammatory markers (**Fig. 6e, f**; **Fig. S9d, e**).

Furthermore, at tumor borders, LAMP1, a lysosome marker, was upregulated in DKO tumors (**Fig. 6g**). The extent of myeloid activation appeared overall stronger in *PB2/PD1* DKO than *PD1* single KO tumors (compare Fig. S5).

### Plexin loss exposes cellular debris to TAMs and reshapes myeloid cell programs from immunosuppression to activation

We next analyzed activation states of TAMs (tumor associated microglia/macrophages) by taking advantage of our snRNA-seq data of *PD1* KO vs. control xenografts (**Fig. 7a**). TAMs from *PD1* KO tumors showed upregulation of immune activation genes such as *SPP1* and *Myo1f* (expressed in macrophages and neutrophils; regulation of adhesion, migration and immune response to pathogens ^30^) (**Fig. 7b**). *Pdgfb,* which has been associated with increased immune infiltration ^31^, was also upregulated in TAMs of KO tumors. In contrast, immune suppression genes were downregulated in *PD1* KO tumors, including *Mrc1*/*CD206* (an anti-inflammatory marker), *Kdm6b* (epigenetic factor promoting immunosuppression in GBM ^32^), and *Trafd1* (TRAF-type zinc finger domain containing 1; immune regulator that primarily suppresses immune responses) (**Fig. 7c**). When analyzing the broader set of DEGs of TAMs in *PD1* KO vs. control tumors, GO terms featured lymphocyte activation and positive regulation of motility (**Fig. 7d, e**).

**Figure 7.**
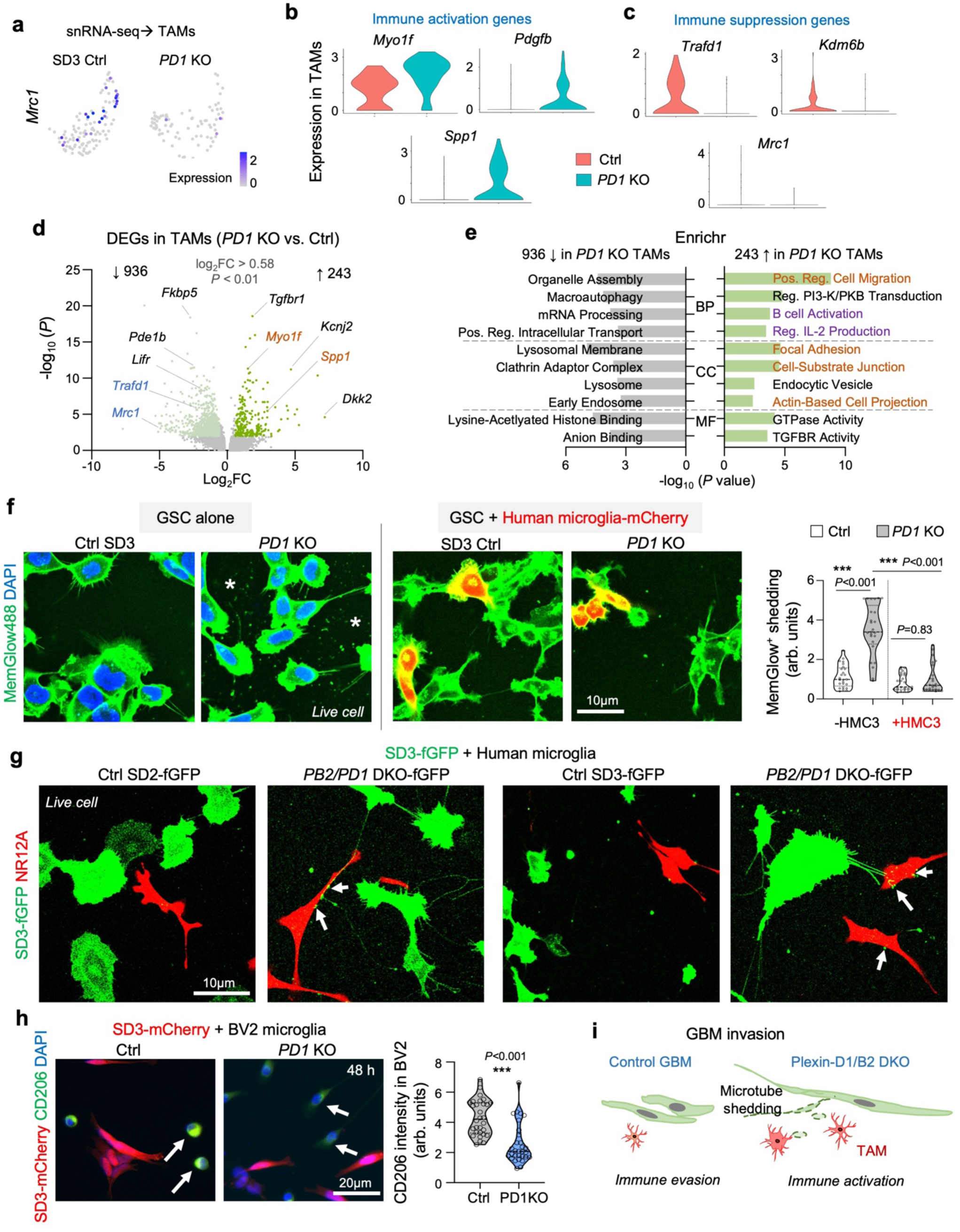
Immune activation in response to tumor invasion in Plexin-D1/B2 double deletion GBM. **a)** Feature plots show expression of immune suppression marker *Mrc1* in TAMs of control and *PD1* KO SD3 GBM. **b, c)** Violin plots show differential expression of immune activation (b) and suppression (c) genes in TAMs of *PD1* KO vs. control SD3 GBM. **d)** Volcano plot showing DEGs in TAMs of *PD1* KO vs. control SD3 GBM. **e)** Enrichr analysis of TAM DEGs of *PD1* KO vs. control SD3 GBMs. BP, biological processes; CC, cellular component; MF, molecular function. **f)** Left, IF images show MemGlow^+^ fragments from cultured *PD1* KO cells (asterisks). Right, co-cultures of SD3 GSCs with human microglia expressing mCherry. Note reduction of MemGlow^+^ fragments in co-culture study with microglia. Quantifications from n=25 regions from 5 independent cultures for each group. Violin plots show median and quartiles. Two-way ANOVA with Tukey’s multiple comparisons test. **g)** Frames from live-cell imaging of fGFP^+^ SD2 (left) and SD3 (right) GBM cells co-cultured with human microglia (HMC3). Note formation of long processes from *PB2/PD1* DKO cells. Also note fGFP^+^ puncta phagocytosed by neighboring cells labeled by membrane dye NR12A (arrows). **h)** IF of co-cultures of mCherry^+^ SD3 GSCs with BV2 microglia show lower expression of immunosuppression marker CD206 (arrow) in microglia in co-culture with *PB2/PD1* DKO GSCs compared to cultures with control GSCs. n=30 cells from 3 independent cultures for each group. Violin plots show median and quartiles. Unpaired two-tailed Student’s t-test. **i)** Model depicting immune-silent migration of control GBM cells, while *PLXND1/B2* double deletion not only impairs invasion and TM retraction but also triggers myeloid cell activation.

To further understand microglial activation in response to cell fragments shed from *PD1* KO cells, we conducted live-cell imaging of GSCs cultured alone or together with mCherry expressing human microglia. In cultures containing only GSCs, we observed the presence of Memglow^+^ fragments, but in co-cultures of GSCs and microglia, MemGlow^+^ fragments released from *PD1* KO GSCs were cleared by microglia (**Fig. 7f**). Similarrly, live-cell imaging of *PB2/PD1* DKO GSCs co-cultured with microglia revealed that fGFP-labeled GSC fragments were sheared off and were taken up by neighboring microglial cells (**Fig. 7g**).

Consistent with the snRNA-seq data, microglia co-cultured with *PD1* KO cells showed downregulation of immune suppression marker CD206 when compared to cultures with control SD3 cells (**Fig. 7h**). Thus, loss of Plexin-D1/B2 resulted in impaired confined migration, fragment shedding, microglial activation, and shifted myeloid programs from immunosuppression to heightened immune activation (**Fig. 7i**).

## DISCUSSION

Diffuse infiltration and immune evasion are defining hallmarks of GBM, leading inevitably to tumor recurrence after initial treatment ^33^. Our study connects these two features of GBM by proposing a unifying mechanism, an increased mechanocompliance of GBM cells via Plexin-D1/B2 to achieve immune-silent invasion.

Invading GBM cells need to detach from the tumor bulk, shift from stationary to invasive state, and penetrate constricted interstitial space without membrane rupture or shedding of immunogenic debris. Our data support a Plexin-mediated cell-state switch, from stationary/connected condition in GBM core to an invasive/disconnected state at GBM border, involving disentanglement from the TM network by enhancing membrane turnover and cytoskeletal dynamics. Loss of either Plexin receptor compromises these functions; loss of both results in increased TM outgrowth, destabilization of plasma membrane, and shedding of cellular debris, thereby impairing confined migration and triggering myeloid activation. Correspondingly, single-nucleus RNA-seq revealed depletion of invasive/mesenchymal-like tumor states and reprogramming of tumor-associated microglia/macrophages (TAMs) toward activation. Our findings thus provide molecular insight into a recent report that stationary GBM cells that are connected by TM networks migrate more slowly and are inversely correlated with invasion ^6^.

The Plexin-D1 and Plexin-B2 single knockouts exhibited similar in vitro and in vivo invasion defects and comparable transcriptomic and lipidomic changes. They also displayed similar shifts in migratory path selection from axonal tracts toward perivascular routes. This perivascular preference of Plexin-deficient tumors may reflect durotactic bias toward the stiffer basal lamina surrounding brain microvasculature ^34^. Thus, Plexins regulate migration across substrates of distinct stiffness or adhesiveness. Notably, Plexin-D1 was found to be selectively upregulated at the tumor border, whereas

Plexin-B2 is more broadly expressed. Double deletion exacerbated fragment shedding, impaired migration and caused slower proliferation. In vivo, GBM transplants with double deletion showed a marked reduction of invasion and bulk size, highlighting functional redundancy and essential roles of mechanocompliance in GBM progression by preventing cellular stress from membrane rupture and fragment shedding.

A striking phenotype in Plexin-deficient GSCs was the overgrowth of thin, elongated TM-like projections containing F-actin, tubulin, and mitochondria. These structures were observed in GSCs cultured on laminin substrates, in 3D microchannels, and in vivo along invasion routes in the corpus callosum. TMs have been implicated in tumor malignancy and therapy resistance through intercellular communication and cargo trafficking ^35^. Of note, our live-cell imaging did not capture active organelle exchange between neighboring cells through these projections. Together, our live-cell imaging, transcriptomic, and lipidomic data indicate extensive remodeling of membrane composition and trafficking (e.g., lysosome/endocytosis), alongside reorganization of cellular projections and synaptic machinery in Plexin-deficient cells. These changes provide a biochemical foundation for altered mechanical properties and TM withdrawal, consistent with established principles linking endocytic trafficking, adhesion dynamics, and ECM remodeling to cell migration ^36^.

From a signaling perspective, semaphorin binding induces Plexin GAP activity towards Rap/R-Ras GTPases to modulate cell-cell adhesion and actomyosin dynamics ^37^. Plexin-B2 binds class 4 semaphorins, which are broadly expressed in CNS cell types ^38–40^. Plexin-D1, classically activated by Sema3E ^41^, also responds to other semaphorins ^42, 43^. Aberrant Plexin-D1 expression occurs in many cancers to promote angiogenesis and metastasis ^44^. In echo, Sema3E overexpression increases tumor invasion via ErbB2 transactivation ^45^. Our findings add a new mechanism by which Plexin-D1 promotes tumor invasion, by enhancing mechanical resilience during confined migration. Beyond ligand engagement, the ring-shaped Plexin extracellular domain itself may act as a mechanosensor. In endothelial cells, Plexin-D1 senses shear stress via ring deformation independently of semaphorin binding ^23^. Similarly, Plexin-B1/B2 confer mechanosensitivity during epithelial wound repair ^46^ and stem-cell collective behavior ^47^. For confined migration, we recently showed that Plexin-B2 modulates membrane tension through semaphorin independent mechanical pathways via the extracellular ring-shaped domain ^22^. Collectively, these studies position Plexins as multifaceted sensors and regulators of cellular biomechanics. A limitation of our study is the use of patient-derived xenograft (PDX) models in immunodeficient mice; thus, the full spectrum of immune responses to debris from invading GBM cells remains to be explored in immunocompetent systems. Future work is also needed to define ligand dependencies and determine whether Plexin-mediated mechanocompliance operates independently of semaphorin binding.

In summary, our findings uncover a mechanical vulnerability of invading GBM cells. Plexin-D1/B2 protect infiltrative tumor cells from membrane rupture and shedding of immunogenic cell fragments during confined migration. Therapeutically targeting Plexins using function-blocking agents, such as antibodies or cross-linking peptides ^48–51^, could force invasive GBM cells into a mechanically fragile, debris-shedding state that may simultaneously curb GBM infiltration and enhance immune recognition, potentially synergizing with immunotherapies.

## METHODS

### Mice

All animal procedures were performed in accordance with the protocols approved by the Institutional Animal Care and Use Committee (IACUC) of Icahn School of Medicine at Mount Sinai. The animal facility light cycle runs for 12 hours daily (7 am to 7 pm), and ambient temperature is 18-23°C with 40-60% humidity. Adult immunocompromised ICR-SCID mice (IcrTac:ICR-Prkdc^scid^) were purchased from Taconic Biosciences for intracranial transplant studies.

### GBM patient data and tissue analysis

Expression and survival analysis of GBM patients were performed with the GlioVis analysis platform ^52^ using glioma datasets from the Cancer Genome Atlas (TCGA ^53^) and the Chinese Glioma Gene Atlas (CGGA ^54^), as well as the Rembrandt data set ^55^. Spatial transcriptomics analysis of Visium 10x datasets of GBM patients ^25^ was performed with the Seurat platform. Paraffin-embedded and formalin fixed (FFPE) sections of GBM patients were obtained from the Mount Sinai Neuropathology core.

### Human glioma cell lines

Human glioma stem cell lines (SD1-4) were originally established from GBM patients at the University of California, San Diego ^56^ and maintained under serum-free human neural stem cell media (HNS) containing Neurocult NS-A basal medium, Neurocult human NS-A proliferation supplement (Stemcell Technologies), 0.0002% heparin, 10 ng/ml bFGF (Peprotech), and 20 ng/ml EGF (Peprotech) as adherent cultures on laminin-coated dishes. Tissue culture dishes were coated with laminin (Invitrogen, 10 µg/ml in PBS) for 1 h at 37°C. Cells were passaged by dissociation with Accutase (Gibco), incubated for 3 min in 37°C incubator before dissociation with a micropipette. Two volumes of basal NS-A media were added to dilute Accutase, and cells were pelleted by centrifugation at 300 g in a tabletop centrifuge (Eppendorf 5702) for 3 min and then resuspended in HNS media before plating.

### Lentiviral CRISPR/Cas KO of Plexins

The plasmid pLentiCRISPRv2 ^57^ was modified by insertion of single guide (sg) RNA sequences targeting human *PLXNB2* or *PLXND1*, or *GFP* as negative control. For *PLXNB2*/*D1* double KO, a lentivirus plasmid was obtained encoding Cas9 and two independent sgRNAs (Vectorbuilder). All lentiviral vectors also express puromycin resistance gene.

sgRNA guide sequence for *PLXNB2* single KO: GTTCTCGGCGGCGACCGTCA (PAM: CGG), deposited as Addgene plasmid 86152. For *PLXND1* single KO: TCTTCCTGGGCACGGTCAAC (PAM: GGG), deposited as Addgene plasmid 178455. For GFP control KO: GGGCGAGGAGCTGTTCACCG, deposited as Addgene plasmid 86153.

sgRNA guide sequences in double KO vectors: For DKO in SD2 cells, for *PLXNB2* TGCGGCTGGTGCGTCGTCGA (PAM: GGG) and for *PLXND1* GCTGCCGTAACCGGTGTACG (PAM: TGG); VectorBuilder ID: VB221008-1210aze. For DKO in SD3 cells: for *PLXNB2* GTTCTCGGCGGCGACCGTCA (PAM: CGG) and for *PLXND1* GCTGCCGTAACCGGTGTACG (PAM: TGG); VectorBuilder ID: VB230501-1188udg.

Lentiviral particles were generated by co-transfection of 293T cells with lentivirus plasmid, the envelope plasmid pMD2.G, and the packaging plasmid psPAX2 (Addgene 12259 and 12260; deposited by Didier Trono, EPFL Lausanne). GSC cultures were infected overnight with lentiviral particles and polybrene (8 μg/ml; Sigma) in six-well plates (2 × 10^5^ cells per well), and after two days, transduced cells were selected with 1 μg/ml puromycin in media.

### Microchannel migration assays

The passage of GBM cells through narrow pores was studied with microchannel devices consisting of polydimethylsiloxane (PDMS) polymer structures on 35 mm glass bottom dishes with parallel rows of microchannels (10 µm height, 12 µm width) containing constrictions of 3 or 8 µm (4D Cell, #MC011). Before seeding of cells, the microchannels were coated with laminin (100 µg/ml in PBS; Invitrogen) for 1 hour at 37°C. Entry ports of microchannel cultures were seeded with 3x10^4^ - 10^5^ GSCs, and after incubation for 4 - 24 h, cells were labeled with dyes MemGlow 488 (Cytoskeleton; 1:200), SPY555-Actin (Cytoskeleton; 1:1000), and NucSpot Live 650 (Biotium; 1:1000). Imaging of migrating GSCs in the microchannels was performed every 5 min on a LSM 780 (Zeiss) confocal microscope (heated stage, 37°C, 5% CO_2_) for up to 17 h. Image analysis was performed with ImageJ (MTrackJ plugin) to determine velocity of cells through constrictions by tracking cells from the time the front of the cell reached a constriction until rear of cell completely exited constriction. Stalling times were measured as the time a migrating cell would remain stalled in front of a constriction before entering it.

### Intracranial transplantation of tumor cells

Adult ICR-SCID mice were used for transplantation studies. Mice were anaesthetized in an induction chamber with a 2.5% isoflurane/oxygen mixture and secured to a stereotaxic apparatus (Stoelting). Anesthesia was maintained with a 1.5% isoflurane/oxygen mixture delivered via a nosecone. After application of lubricating ophthalmic ointment (Artificial Tears, Akron), a cranial burr hole was drilled through a scalp incision at coordinates of 2.0 mm lateral and 0.5 mm anterior to bregma. Human GBM cells (1x10^5^) suspended in 5 µl PBS were injected through the trephination at a depth of 3.2 mm using a 10 µl gas-tight syringe (Hamilton) linked to a Nanomite programmable syringe pump (Harvard Apparatus) with a constant infusion rate of 1 µl/min to prevent backflow. After injection, the syringe was incrementally raised using the stereotaxic apparatus over 5 min period. Scalp incision was sealed using a tissue adhesive (Vetbond, 3M). Tumor bearing mice were euthanized for collection of brain tissue after a specified time.

### Immunofluorescence staining and immunohistochemistry

Mice carrying intracranial GBM transplants were intracardially perfused with ice-cold PBS followed by ice-cold 4% PFA/PBS. Brains were harvested and fixed overnight in 4% PFA/PBS followed by two successive overnight incubations in 12.5% and 25% sucrose/PBS. Brains were then embedded in O.C.T compound (Fisher Scientific) and frozen on dry ice. Cryosections of 20 µm thickness were cut using a cryostat (Leica) and collected as floating sections in anti-freeze media (25% glycerol, 30% ethylene glycol and 0.05 M phosphate buffer) and stored at -20°C.

For immunofluorescence (IF) staining, floating sections were blocked for 1 h (blocking buffer: PBS with 5% donkey serum and 0.3% Triton X-100), then incubated overnight with primary antibodies in antibody dilution buffer (PBS with 1% BSA and 0.3% Triton X-100). This was followed by staining with Alexa-labeled secondary antibodies (Jackson ImmunoResearch) for 2 h, and counterstaining with DAPI (Invitrogen). Sections were washed in PBS and mounted on glass slides with Fluoromount G (Southern Biotech).

For IF of FFPE patient specimens, microtome sections were de-paraffinized with Histo-clear (National Diagnostics, HS2001GLL), followed by dehydration with 100%, 95%, and 80% ethanol (Fisher Scientific, BP28184). Antigen retrieval was performed with Visucyte Antigen Retrieval Reagent-Basic (R&D Systems, VCTS021) at 95°C for 30 minutes, followed by cooling down to room temperature for 20 minutes.

Primary antibodies for IF:

anti-PDGFRα (host: goat), source: R&D Systems AF1062, dilution: 1:200;

anti-Plexin-B2 (sheep), R&D Systems AF5329, 1:200;

anti-Plexin-D1 (goat), R&D Systems AF4160, 1:400;

anti-CD68 (rat), Bio-Rad MCA1957GA, 1:100;

anti-CD16/32 (rat), BD Bioscience 553142, 1:200;

anti-Lamp1 (rat), Developmental Studies Hybridoma Bank 1D4B, 1:200;

anti-CD206 (goat), R&D Systems AF2535, 1:100;

anti-Iba1 (rabbit), Fujifilm Wako 019-19741, 1:500;

anti-Iba1 (goat), Novus Bio NB100-1028, 1:500;

anti-PECAM1 (rat), BD Bioscience 553370, 1:250;

anti-MBP (rabbit), CST 78896, 1:500;

anti-PU.1 (rabbit), eBioscience MA5-15064, 1:200;

anti-human nuclear antigen HNA (mouse), Millipore MAB1281, 1:500;

anti-human NCAM (mouse), Santa Cruz sc-106, 1:200;

anti-EGFR (rabbit), Cell Signaling 4267, 1:250;

anti-integrin β1 (human specific clone TS2/16) (mouse), Santa Cruz sc-53711, 1:100;

anti-pMLC2 (rabbit), Cell Signaling #3671, 1:100;

anti-GAP-43 (mouse, clone B-5): Santa Cruz sc-17790, 1:200.

Secondary antibodies coupled to Alexa fluorophores (Jackson ImmunoResearch) were used in 1:300 dilutions of 1 mg/ml stocks.

Staining for filamentous actin was performed with Alexa Fluor 594-Phalloidin (Invitrogen A12381).

### Western blot

For Western blot protein detection, cells were lysed with RIPA buffer (Sigma) including protease and phosphatase inhibitors (Pierce). Lysates were run on SDS-PAGE gels (4-12% NuPAGE Bis-Tris gradient gels; Invitrogen) and transferred onto nitrocellulose membranes with the Novex gel system (Invitrogen). Membranes were incubated at 4 °C overnight with primary antibodies and then for 1 hour with secondary donkey antibodies IRDye 680 and 800 (Li-Cor). Fluorescent protein bands were detected with the Odyssey Infrared Imaging System (Li-Cor).

Primary antibodies for Western blot analysis: anti-Plexin-D1 (goat), R&D systems AF4160, 1:400; anti-β-actin (host: mouse), Sigma A1978, 1:10,000.

### Image processing and quantification

Images were collected with Zeiss Zen software (Zen Blue v3.6 and Zen Black v8). Image data were quantified with the Fiji Is Just ImageJ (FIJI, v2.16.0) package of ImageJ 75. Data were handled using Microsoft Excel (v16.95.1). To quantify the intensity of immunofluorescence signals, images were spatially calibrated and normalized intensity values were then divided by area of region of interest (ROI). To quantify PU.1^+^ and HNA^+^ cell abundance per unit area, in each ROI selection, the number of PU.1^+^ or HNA^+^ cells was counted manually. Three-dimensional surface rendering of z-stack images of confocal microscopy data was created with the Imaris v10.0 software package (Bitplane).

### Scratch Assay

Cells were seeded in laminin-coated 12-well plates at a density of 1x10^5^ cells per well. After 7 day culture with confirmation of complete confluence of wells, a scratch was made through the center of the dish with 200 µl size micropipette tips (t = 0 h), and wound closure speeds were measured followed by micrographs of identical areas at t = 12, 24, 36 and 48 h after scratch.

### Growth Assay

Cells were seeded in laminin-coated 12-well plates at a density of 1x10^5^ cells per well. After 24 h of incubation in cell culture, cells are dissociated with Accutase (BD Biosciences) and diluted with DPBS (Gibco). After centrifugation with 500 g for 5 min, cell pellets were resuspended in DPBS and cell numbers were counted.

### FluoVolt live staining

To assess membrane potential of cells in 2D cell culture, FluoVolt (Thermo, 1:1000) membrane dye was diluted with Powerload supplement (1:100) in neural stem cell media (Neurocult NS-A proliferation kit (human), Stemcell Technologies). Cells were stained for 30 min at 37 °C, washed twice with PBS and then HEPES (1:100) were added. Images were taken after 15 minutes with 488 nm excitation with an LSM 710 or 780 inverted confocal microscope (Zeiss).

To assess membrane potential of GBM cells, cells were seeded in laminin-coated 4 chamber glass bottom dish (35mm Dish with 20mm Bottom well) at a density of 10^4^ cells per chamber. After 24 h, cells were incubated with FluoVolt dye and imaged as described above.

### GSC-human microglia co-culture and live cell imaging

GSCs (2 × 10^3^ cells) mixed with HMC3 immortalized human microglia (ATCC) ^58^ (2 × 10^3^) were seeded into laminin-coated 4-chamber glass bottom dish (35 mm Dish with 20 mm bottom well) with neural stem cell media. After 2 days of co-cultures, cells were labeled by with NR12A (Cytoskeleton; 1:1000), and NucSpot Live 650 (Biotium; 1:1000) and images were taken every 5 minutes on a LSM 780 (Zeiss) inverted confocal microscope (heated stage, 37 °C, with 5% CO2) for up to 17 h. GSCs were also co-cultured with immortalized mouse microglia line BV2 ^59^ (obtained from laboratory of Dr. Landreth, Indiana University).

### Molecular reporters and probes for GBM cells

A lentiviral vector for expression of membrane anchored farnesylated GFP and neomycin selection cassette was obtained from Vectorbuilder (vector ID: VB240206-1365hmd). A lentiviral vector for constitutive expression of mCherry in GSCs was obtained from Addgene (plasmid 36084; deposited by P. Tsoulfas). Expression plasmids for myrPalm-CFP (Addgene plasmid 14867) and PH-PLCD1-GFP (Addgene plasmid 51407) were obtained from the Addgene repository.

Cells were also stained for live cell imaging with Lysotracker (L7526) or Mitotracker (Invitrogen M7514) membrane dye NR12A (Cytoskeleton MG07) or tubulin markers SPY650-Tubulin (Cytoskeleton CY-SC503).

### Single-nucleus RNA-seq

Nuclei were isolated from brain tissues carrying GSC brain tumor transplants by a lysis protocol optimized to preserve nuclear structure and RNA integrity (#130-128-024, Miltenyi Biotec.). Nuclei were counted, stained with SYBR-green for quality control, and processed with the Evercode Whole Transcriptome v2 kit (Parse Biosciences, Version 2.3-UM0022) to generate cDNA libraries. DNA was sequenced on an Illumina NovaSeq X device using a 25B flow cell, generating paired-end 150 bp reads to a depth of ∼50,000 reads per nucleus. Raw read data was processed with Evercode Whole Transcriptome Analysis software (Parse Biosciences) for barcode demultiplexing, alignment to GRCh38 and mouse MM39 reference genomes, and UMI-based quantification, and generation of gene-by-cell count matrices.

Read data was processed using the Seurat (v5.0.1) package ^60^. Clusters were annotated by analysis of canonical marker gene expression. Differential gene expression analysis was performed in Seurat using the FindMarkers function (Wilcoxon rank-sum test) with cutoff of log_2_FC > 0.5. Gene Ontology (GO) enrichment analyses were conducted for upregulated and downregulated genes using the ENRICHR platform (https://maayanlab.cloud/Enrichr/) ^61^. Analyzed gene sets included GO Biological Process 2025, Cellular Component 2025, and Molecular Function 2025. Additional pathway enrichment analysis was performed with WikiPathways (Human, 2024 release) and KEGG (2025 release) gene sets.

### Bulk RNA-seq

Total RNA was isolated from adherent GSC cultures using QIAGEN RNeasy Micro Kit with on-column DNase digest. RNA integrity was assessed on an Agilent 2100 Bioanalyzer, and samples with RIN ≥ 7.0 were advanced. Libraries were prepared with Illumina TruSeq Stranded mRNA (poly(A) selection) reagents, with input ≥ 500 ng total RNA. Libraries were sequenced on an Illumina NovaSeq X device, with parameters paired-end, 2x150 bp, target depth ∼40 M read pairs per sample (∼80 M total reads), and ∼12.08 Gb per sample.

Raw sequencing reads from samples were mapped to hg38 using HISAT2 v2.2.1 using default parameters ^62^. RNA-seq QC metrics were calculated using FastQC v0.11.9 (https://www.bioinformatics.babraham.ac.uk/projects/fastqc/). Counts of reads mapping to genes were obtained using featureCounts software of Subread package v2.0.1 against Ensembl v109 annotation ^63^. Differential expression was done using the DESeq2 v1.42.0 R package ^64^.

### Lipidomics

SD3 GSCs carrying CRISPR/Cas9 KO mutations of the *PLXNB2* and/or *PLXND1* genes were cultured in neural stem cell media in 6-well plates until confluence. For lipidomics analysis, GSCs were dissociated with Accutase, washed with PBS, and pelleted in 1.5 ml Eppendorf tubes. For each sample, pellets of 5-8 x 10^5^ cells were stored at -80°C and subsequently shipped to the Columbia University Biomarkers Core Laboratory for LC/MS targeted lipidomics analysis.

### Statistical analysis

Statistical analyses were performed using GraphPad Prism 10 software, using the setting NEJM (New England Journal of Medicine) for reporting of *P* values. *P* < 0.05 was considered as statistically significant and *P* values < 0.001 are grouped as such. Bar graphs represent the means and error bars represent the s.e.m. Data distribution was assumed to be normal, but this was not formally tested. A one-way or two-way analysis of variance (ANOVA; for three or more experimental groups) and Student’s t-test were performed to assess whether experimental groups were significantly different from each other. No data were excluded from statistical analysis. Data collection experiments were not randomized, and investigators were not blinded.

## Supporting information

Supplementary figures with legens

Legends for videos and tables

## Data availability

The RNA-seq data generated in this study were deposited at the NCBI Gene Expression Omnibus (GEO) database under accession numbers GSExxxxxx (for single nucleus RNA-seq of intracranial transplant) and GSExxxxxx (for RNA-seq of cultured SD2 and SD3 cells carrying mutations of *PLXNB2* and/or *PLXND1*). All data supporting the findings of this study are available from the corresponding authors upon reasonable request. Source data are provided with this paper.

## Acknowledgements

We thank the Columbia University Biomarkers Core Laboratory for performing lipidomics studies.

S.K. was supported by an NIH/NCI T32 training award (T32CA078207, to Tisch Cancer Institute at Icahn School of Medicine at Mount Sinai) and C.J.A. by an NIH/NINDS career development award (K01NS127948). This study was also supported by research grants from the NIH/NINDS to H.Z. (R01NS134159), to H.Z. and C.J.A. (R21NS145550), to R.H.F. (R01NS092735, R21NS134158), and to N.M.T. (R01NS106229).

## Author contributions

S.K., C.J.A., R.H.F., and H.Z. conceptualized the study, planned experiments, and analyzed data. S.K., C.J.A., H.T., C.K., S.S., and A.D. carried out cell assays. S.K. and J.Z. performed xenograft studies. S.K., C.J.A., J.S.F, A.R., and L.S. processed and analyzed RNA-seq data. N.M.T. supported human tissue analyses. R.H.F. and H.Z. wrote the manuscript.

