## Supplementary figures with legens for "Guidance receptor-mediated mechanocompliance of GBM cells facilitates immune-silent invasion"

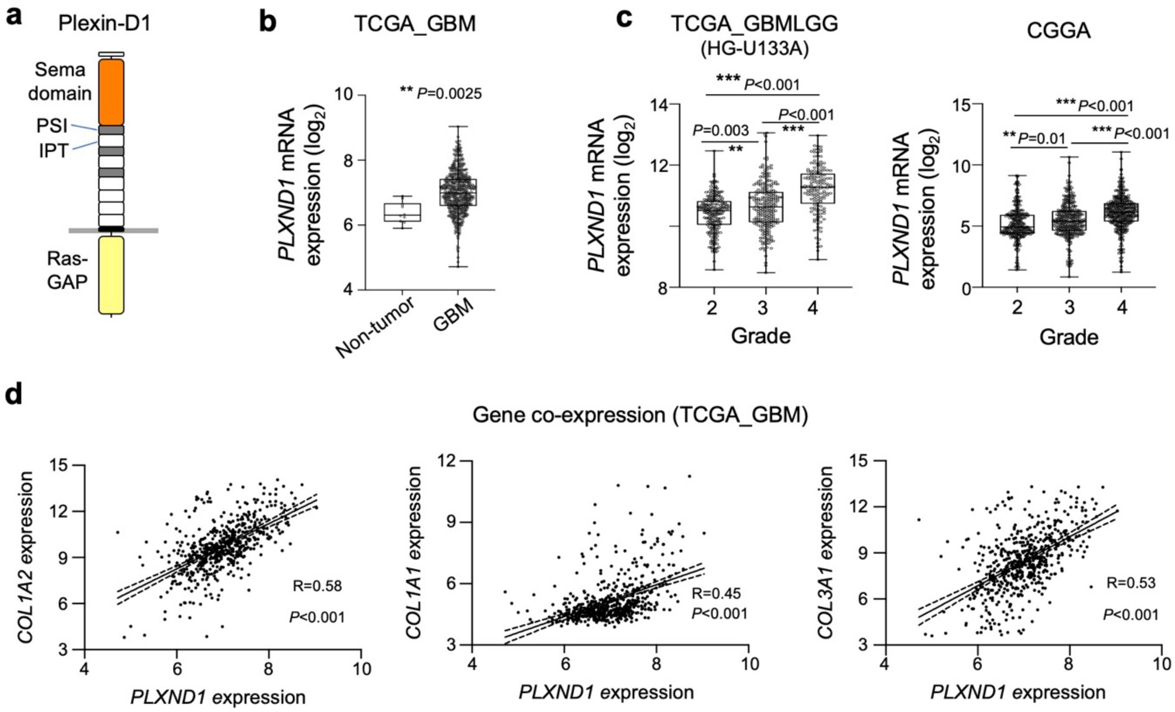


**Figure S1. Plexin-D1 expression in high-grade gliomas.**

**a)** Diagram of Plexin-D1 domains.

**b)** Elevated expression of *PLXND1* in GBM tumors of TCGA dataset (at GlioVis platform).

**c)** *PLXND1* expression increases with glioma grade in mixed high-grade and low-grade gliomas, datasets of TCGA and CGGA patient databases at GlioVis platform.

**d)** Co-expression of *PLXND1* with collagen genes, TCGA_GBM patient dataset.

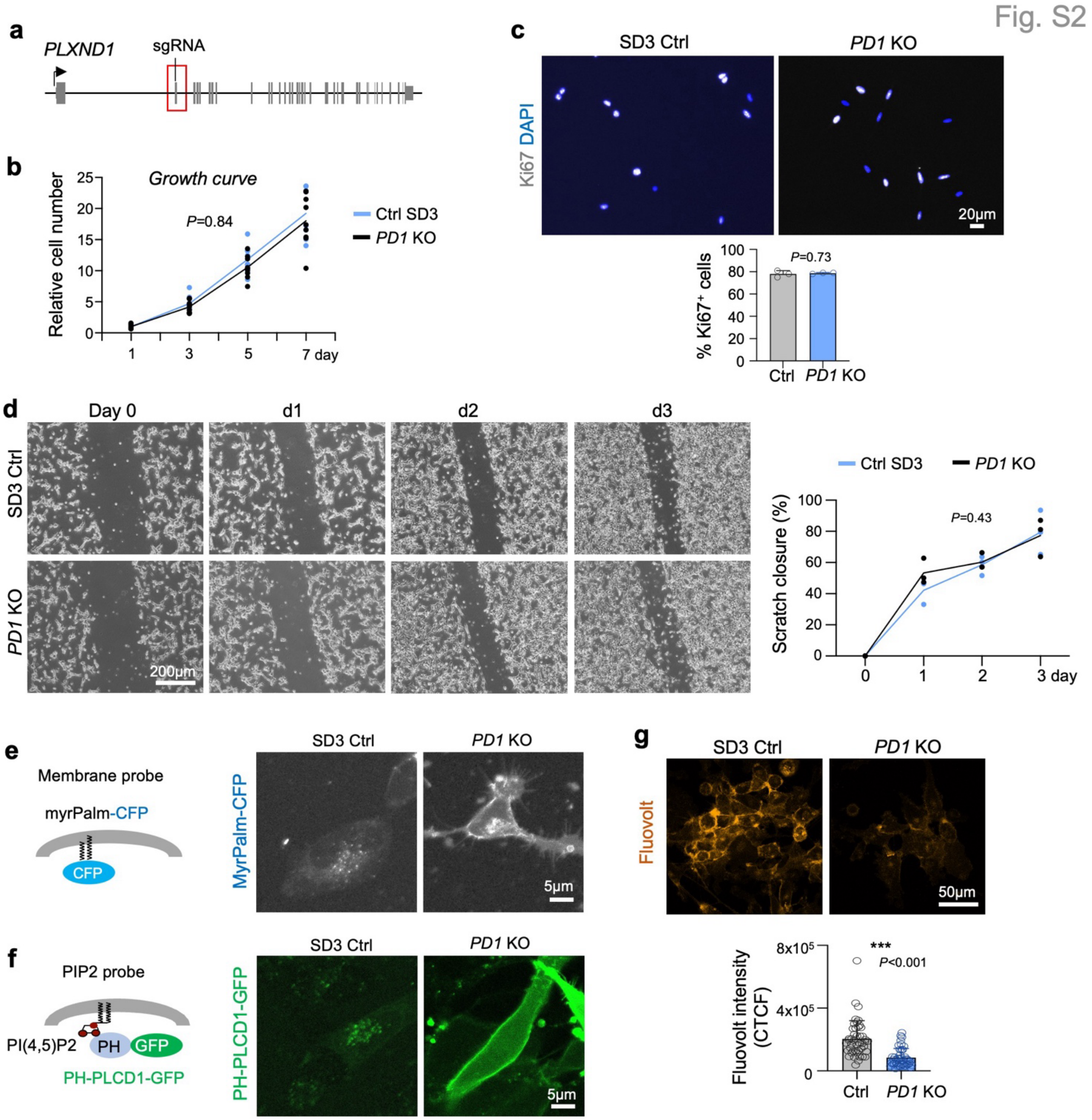


**Figure S2. Plexin-D1 KO reduces membrane turnover and alters membrane electric charge**

**a)** Cas9/sgRNA targeting of second coding exon of human *PLXND1* gene.

**b)** Growth curve study shows no significant effect of *PLXND1* knockout (*PD1* KO) on proliferation of SD3 GBM cells. Two-away ANOVA, n=9 independent cultures for each condition. Data curve represents means values.

**c**) No significant difference of proliferation marker Ki67 expression between SD3 control cells *PD1* KO GSCs. n=3 independent cultures. Unpaired two-tailed Student's t-test. Bar graphs represent mean ± s.e.m.

**d**) Images and quantification of scratch wound closure assay. n=3 independent cultures for each condition. Two-away ANOVA. Data curve represents means values.

**e**) Left, diagram of myr-Palm-CFP fluorescent probes that is anchored to inner leaflet of plasma membrane. Right, representative images show internalization of probe in control SD3 cells, but retainment at plasma membrane in *PD1* KO cells at 72 h post-transfection.

**f**) Schematic of PH(PLCD1)-GFP PIP2 probe. Right, live-cell imaging at 72 h post transfection reveals that the PH(PLCD1)-GFP probe was internalized in control GSCs but retained on membrane of *PD1* KO GSCs.

**g**) Fluorescence imaging shows reduced signals of voltage-sensitive membrane dye Fluovolt in *PD1* KO cells compared to controls. n=45 cells. Bar graphs show mean ± s.e.m. Unpaired two-tailed Student's t-test. CTCF, corrected total cell fluorescence.

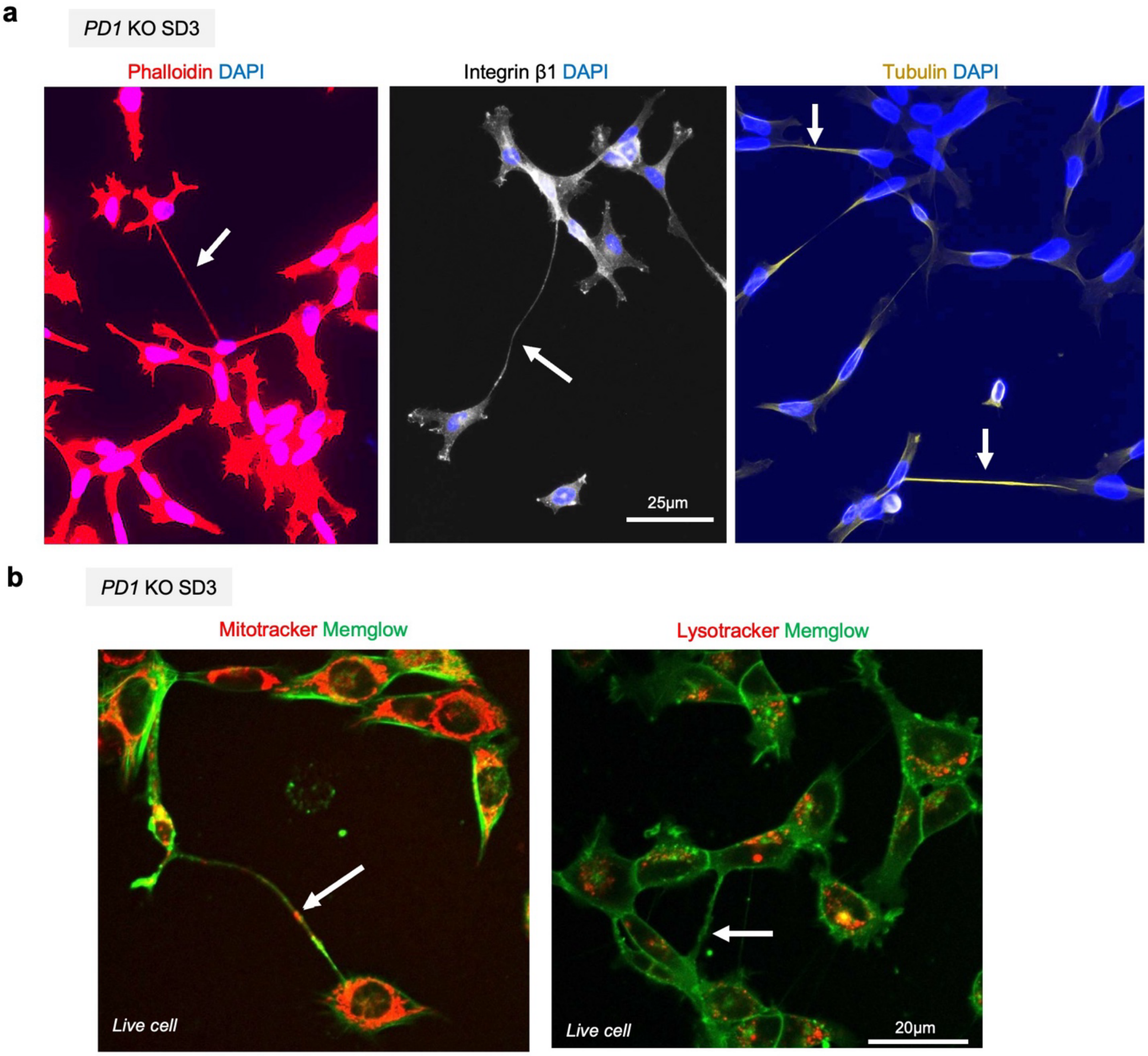


**Figure S3. Plexin-D1 KO cells extend tumor microtube-like processes.**

**a)** Fluorescence images reveal thin long TM-like processes (arrow) containing F-actin, tubulin, and integrin β1.

**b)** Live-cell images show the presence of mitochondria (arrow) in the processes projecting from SD3 *PD1* KO cells. In contrast, lysosome content in TM-like processes was low.

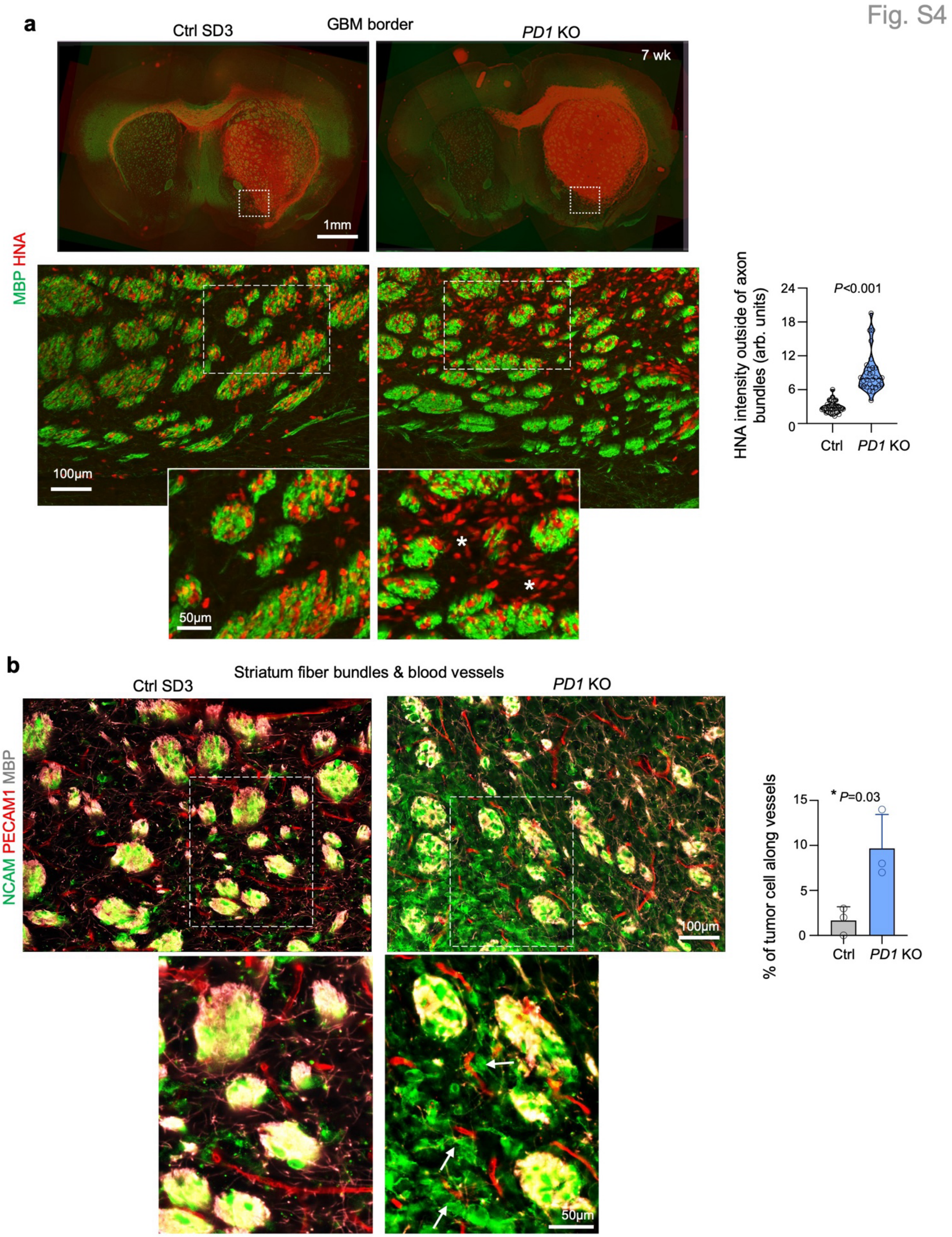


**Figure S4. Plexin-D1 KO shifts preferences for migration paths in GBM invasion.**

**a)** IF overview images and enlarged views show relationship of invading GBM cells ($HNA^+$) and striatal axon fiber bundles ($MBP^+$) at GBM borders. In control SD3 transplants, tumors showed preference for fiber tracks, whereas in *PD1* KO tumors, tumor cells also invade in between axon bundles (asterisks). Quantifications from n=30 regions from three tumors for each condition. Unpaired two-tailed Student's t-test. Violin plots show median and quartiles.

**b)** IF images with enlarged views show juxtaposition of invading *PD1* KO tumor cells to blood vessels in-between axon fiber bundles (arrows). Quantifications from n=3 independent transplants for each condition. Unpaired two-tailed Student's t-test. Bar graphs show mean ± s.e.m.

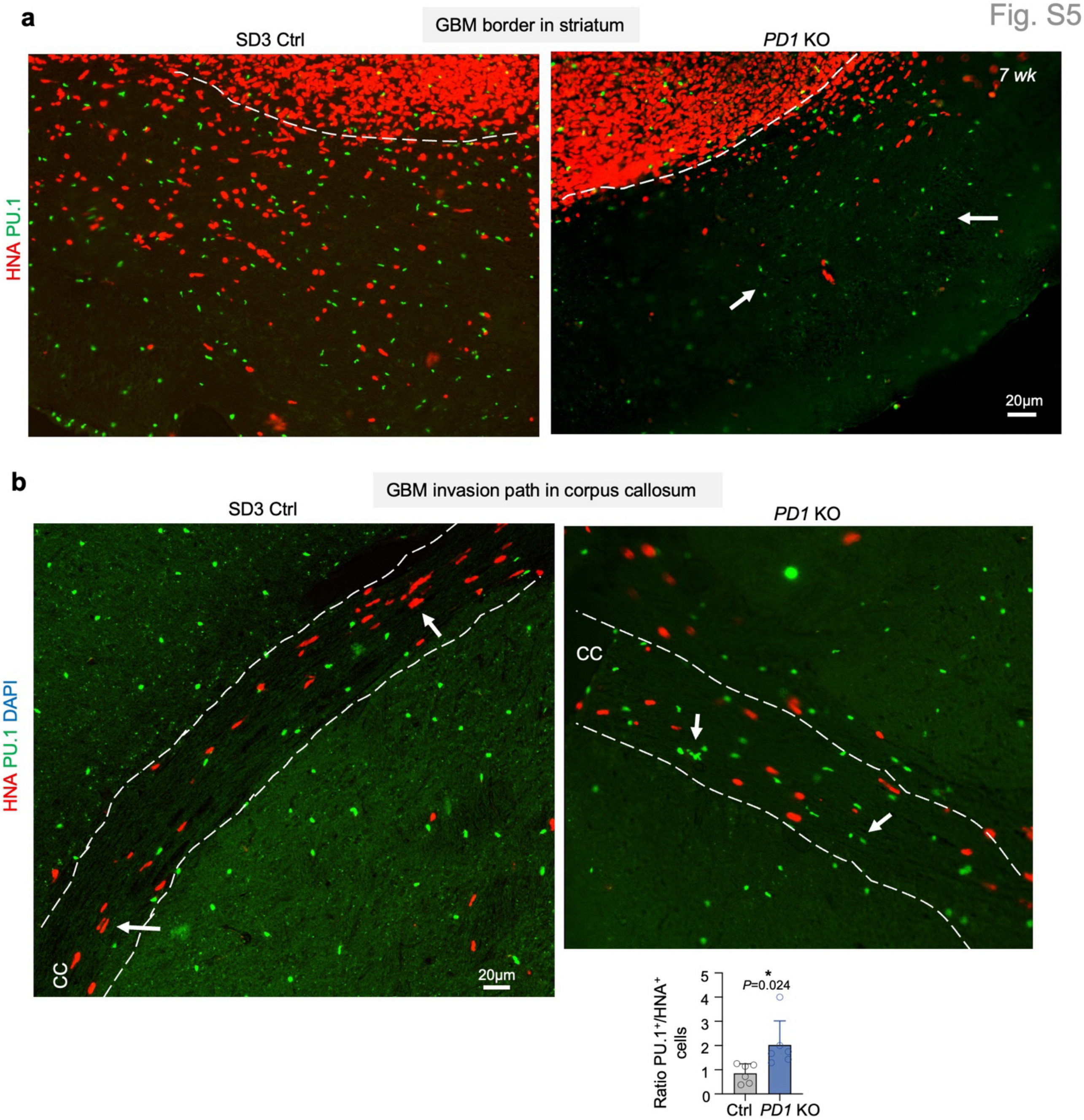


**Figure S5. Increase of myeloid cells at GBM border and in invasion paths in Plexin-D1 KO tumors.**

**a)** Representative IF images show reduced infiltration of Plexin-D1 KO GBM tumors at border compared control tumors. Note abundant PU.1$^+$ myeloid cells at GBM border (arrows) despite reduced invading tumor cells in *PD1* KO.

**b)** IF and quantification show increased ratio of PU.1$^+$ myeloid cells to HNA$^+$ GBM cells along invasion path of corpus callosum (CC). Quantifications from n=6 CC regions from three tumors for each condition. Unpaired two-tailed Student's t-test. Bar graphs show mean ± s.e.m.

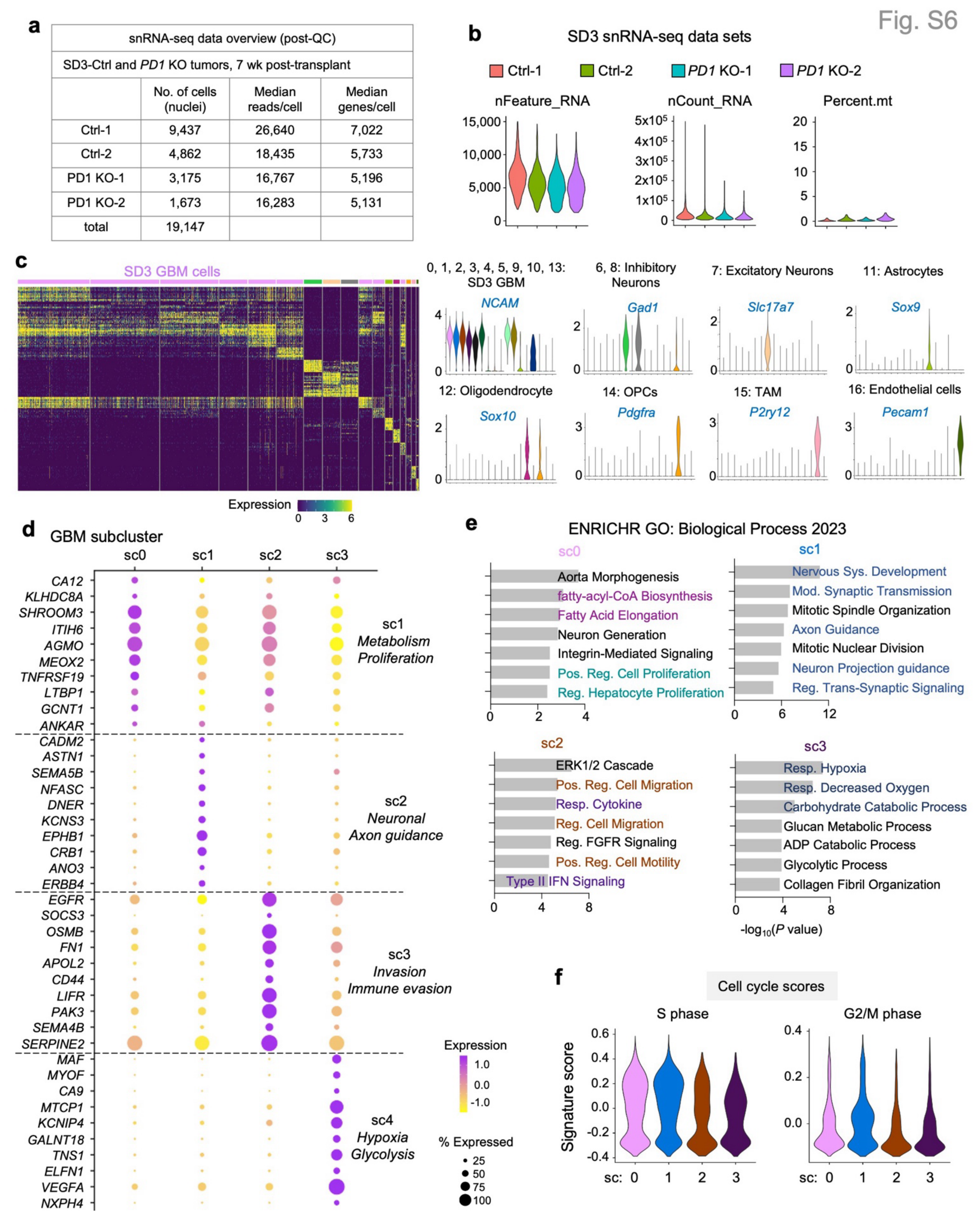


**Figure S6. Single nucleus RNA-seq reveal distinct cell states of bulk growth vs. infiltrate GBM cells in SD3 tumor**

**a)** Table listing number of high-quality nuclei sequenced, with n = 2 for each genotype.

**b)** Violin plots showing quality metrics of snRNA-seq samples.

**c)** Heatmap showing distinct gene expression of tumor and stromal cell clusters in SD3 tumors. Right, violin plots show marker gene expression for each major cluster

**d)** Dot plots of top DEGs for each subcluster of GBM cells in SD3 tumor. Right, main function annotations.

**e)** Enrichr pathway analysis of marker genes for each tumor cell subcluster in SD3 tumor, color coded for distinct functional pathways.

**f)** Violin plots show cell cycle across four subclusters of tumor cells, showing no overt difference of S phase or G2M score.

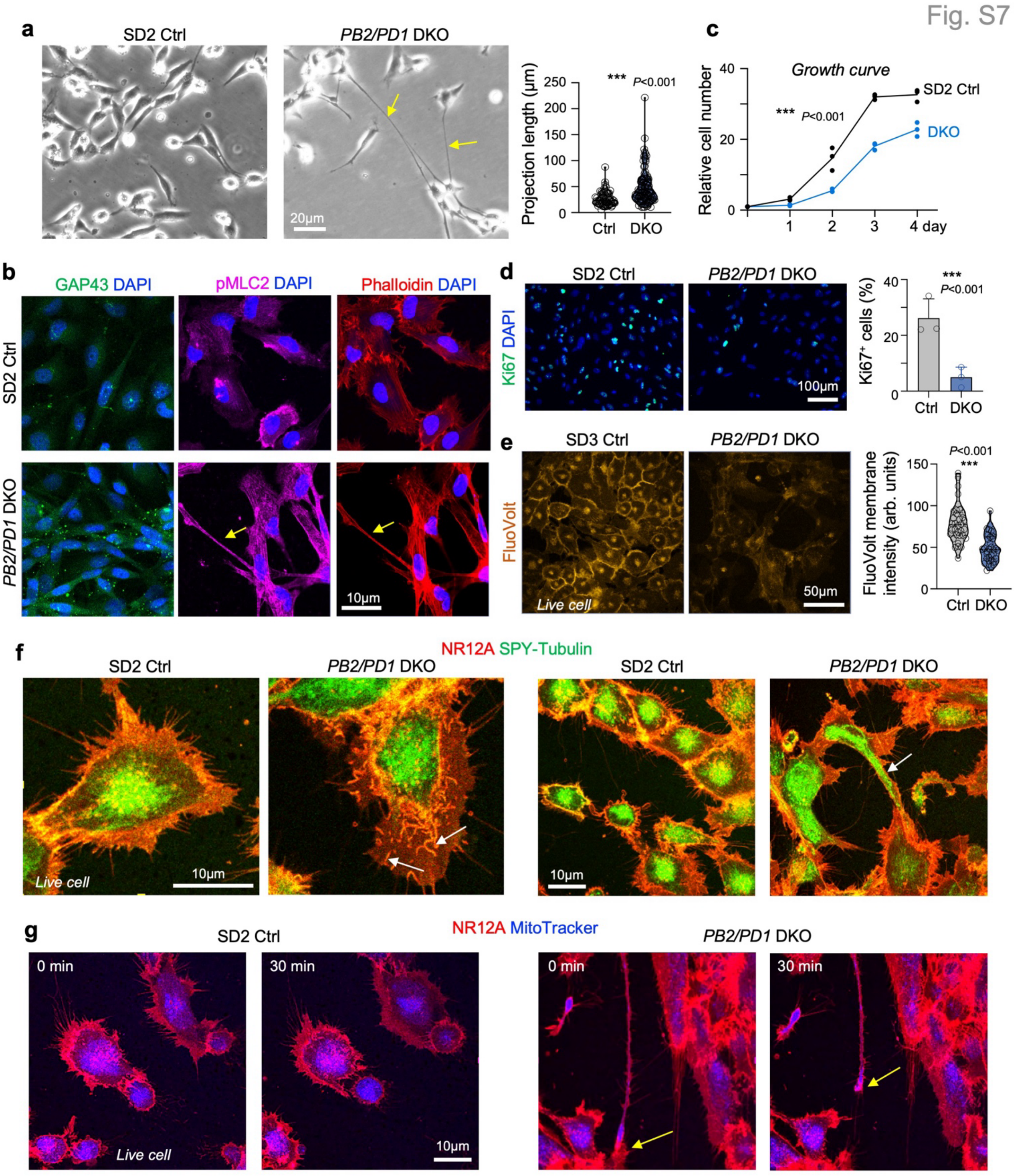


**Figure S7. Double deletion of Plexin-D1/B2 results in TM overgrowth and GAP43 upregulation.**

**a)** Phase contrast images show long thin processes (arrows) from cultured SD2 *PB2/PD1* DKO GSCs. Quantification from n=120 control cells and n=60 *PB2/PD1* DKO cells from three independent experiments. Violin plots show median and quartiles. Unpaired two-tailed Student's t-test.

**b)** IF staining shows increased GAP43 expression in SD2 *PB2/PD1* DKO cells compared to control cells. The TM-like processes contained pMLC2 and F-actin (arrows).

**c)** Growth curve analysis shows significant lower proliferation of SD2 *PB2/PD1* DKO cells. Two-away ANOVA, n=3 independent cultures.

**d)** IF staining shows reduced proportion of $Ki67^+$ SD2 *PB2/PD1* DKO cells compared to control GSCs. Bar graphs represent mean ± s.e.m. Unpaired two-tailed Student's t-test. n=3 independent cultures.

**e)** Reduced Fluovolt signals in SD3 *PB2/PD1* DKO cells. Quantification from n=34 cells for each condition. Violin plots show median and quartiles. Unpaired two-tailed Student's t-test.

**f)** Representative images captured from time-lapse imaging show examples of increased membrane ruffles (arrows in left images) as revealed by membrane dye NR12A. Note presence of tubulin in TM (arrows in right images). Related to Videos S3 and S4.

**g)** Representative images captured from time-lapse imaging show withdrawal of the tip of TM with abundant mitochondria (arrow). Related to Video S5.

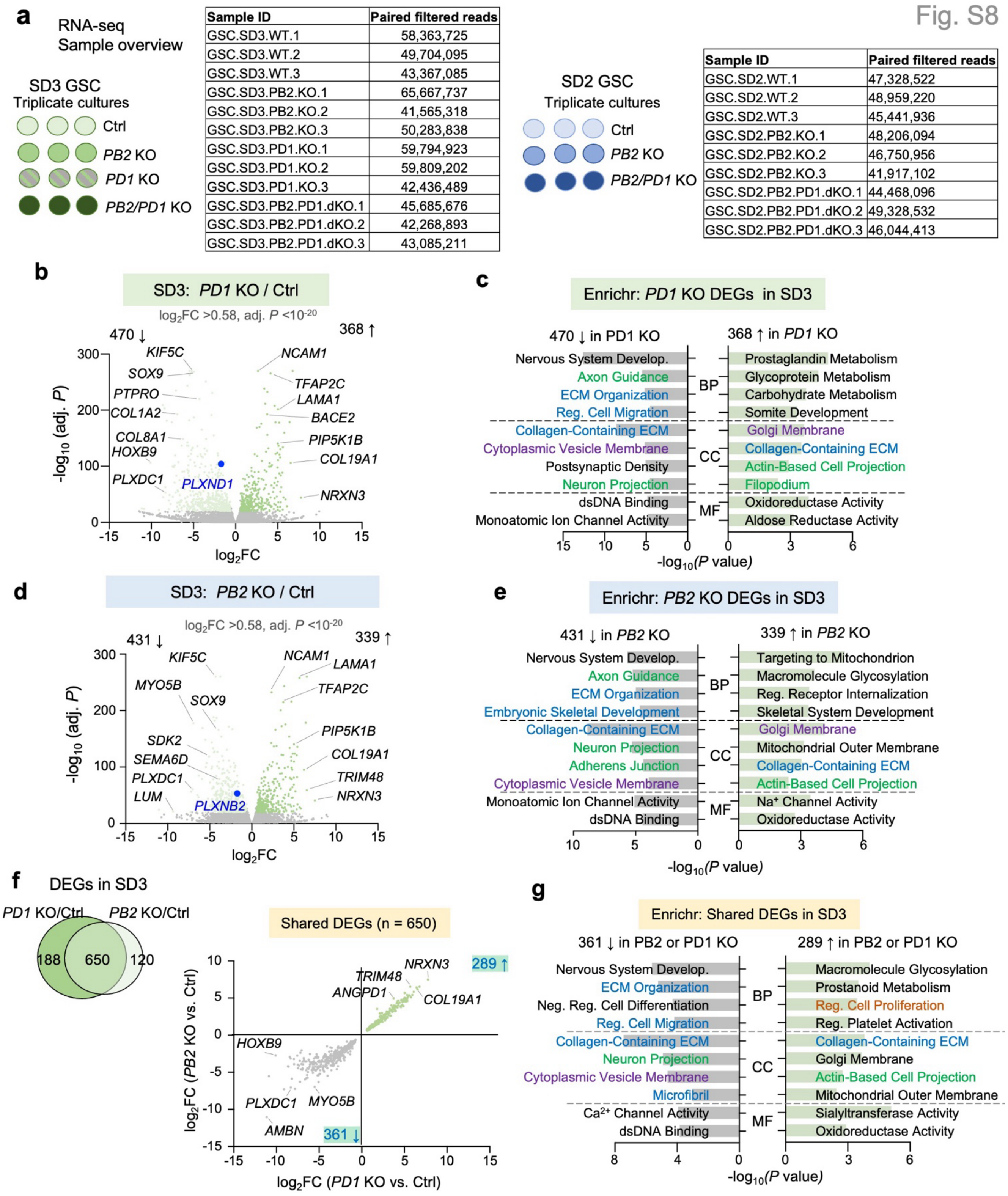


| Sample ID | Paired filtered reads |
|---|---|
| GSC.SD3.WT.1 | 58,363,725 |
| GSC.SD3.WT.2 | 49,704,095 |
| GSC.SD3.WT.3 | 43,367,085 |
| GSC.SD3.PB2.KO.1 | 65,667,737 |
| GSC.SD3.PB2.KO.2 | 41,565,318 |
| GSC.SD3.PB2.KO.3 | 50,283,838 |
| GSC.SD3.PD1.KO.1 | 59,794,923 |
| GSC.SD3.PD1.KO.2 | 59,809,202 |
| GSC.SD3.PD1.KO.3 | 42,436,489 |
| GSC.SD3.PB2.PD1.dKO.1 | 45,685,676 |
| GSC.SD3.PB2.PD1.dKO.2 | 42,268,893 |
| GSC.SD3.PB2.PD1.dKO.3 | 43,085,211 |

| Sample ID | Paired filtered reads |
|---|---|
| GSC.SD2.WT.1 | 47,328,522 |
| GSC.SD2.WT.2 | 48,959,220 |
| GSC.SD2.WT.3 | 45,441,936 |
| GSC.SD2.PB2.KO.1 | 48,206,094 |
| GSC.SD2.PB2.KO.2 | 46,750,956 |
| GSC.SD2.PB2.KO.3 | 41,917,102 |
| GSC.SD2.PB2.PD1.dKO.1 | 44,468,096 |
| GSC.SD2.PB2.PD1.dKO.2 | 49,328,532 |
| GSC.SD2.PB2.PD1.dKO.3 | 46,044,413 |

**Figure S8. RNA-seq shows convergent transcriptional impact by Plexin-D1 and -B2 deletion in SD3 GSC.**

**a)** Table listing triplicate samples of SD3 or SD2 GSCs with single or double deletion of Plexins D1/B2.

**b)** Volcano plots showing DEGs between SD3 *PD1* KO and control cells, with *PLXND1* highlighted as downregulated DEG.

**c)** Enrichr analysis of the up and down-regulated DEGs (*PD1* KO vs. control) in SD3 GSCs, color-coded for distinct functional pathways.

**d)** Volcano plot shows the DEGs of SD3 *PB2* KO vs. control cells, with *PLXNB2* highlighted as downregulated DEG.

**e)** Enrichr analysis of SD3 *PB2* KO vs. control up and down-regulated DEGs, color-coded for distinct functional pathways.

**f)** Left, Venn diagram shows overlap of SD3 *PD1* KO and *PB2* KO DEGs. Right, graphs show converging directionality of shared *PD1* KO and *PB2* KO DEGs.

**g)** Enrichr analysis of shared up and down-regulated DEGs (SD3 *PB2* KO vs. control and SD3 *PD1* KO vs. control), color-coded for distinct functional pathways.

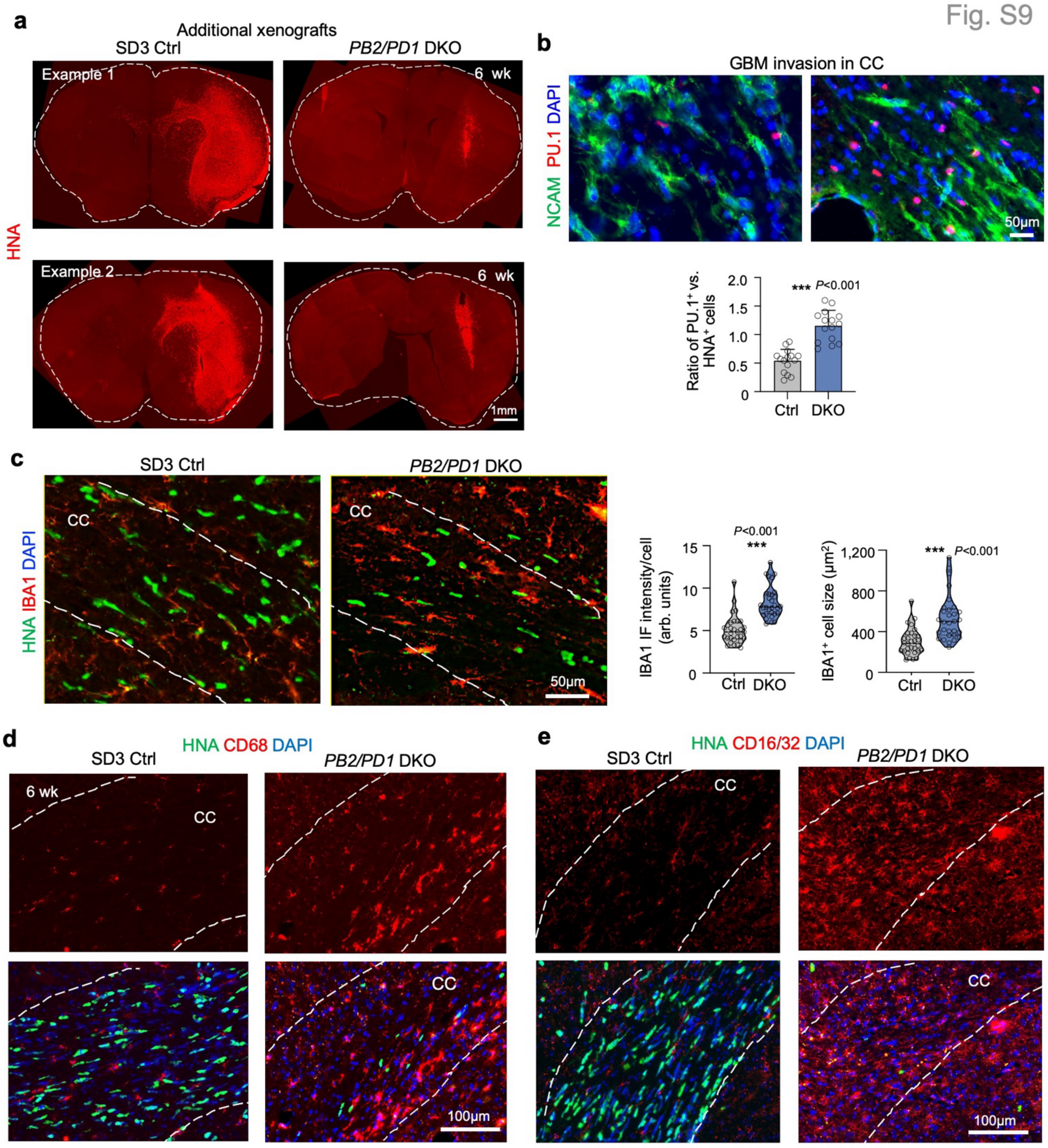


**Figure S9. Enhanced immune recruitment and activation in GBM tumors with double deletion of Plexin-B2/D1.**

**a)** Additional examples of expansion of SD3 control and *PB2/PD1* DKO tumors, visualized by IF for human nuclear antigen (HNA) at 6 weeks post-transplant.

**b, c)** IF images show that in SD3 *PB2/PD1* DKO tumors, invading GBM cells in corpus callosum (CC) were surrounded by more PU.1$^+$ and IBA1$^+$ TAMs. For PU.1/NCAM cell ratio quantification, n=15 regions from 3 independent transplants for each group. Bar graphs represent mean ± s.e.m. For IBA1 quantification, n=30 cells from 3 independent transplant for each group. Violin plots show median and quartiles. Unpaired two-tailed Student's t-test.

**d, e)** IF images at corpus callosum (CC) show higher signal for phagocytosis marker CD68 (d) and pro-inflammatory marker CD16/32 (e) in SD3 *PB2/PD1* DKO tumors.
