## Supplementary material for "Guidance receptor-mediated mechanocompliance of GBM cells facilitates immune-silent invasion": Legends for videos and tables

**Legends for Supplementary Videos**

**Video S1.** Microchannel assay with SD3 control (left) and *PD1* KO (right) GSCs, revealing reduced migration capacity of *PD1* KO GSCs. Cells were stained with live dyes SPY-actin (red), Memglow (green), and Nucspot (blue).

**Video S2.** Zoom in view of SD3 *PD1* KO GSCs migrating through microchannels. Tethers between cells are visible by SPY-actin staining.

**Video S3.** Live cell imaging of SD2 control (left) and *PB2/PD1* DKO (right) GSCs cultured on laminin coated dishes. Cells were stained with live dyes NR12A (red), SPY-tubulin (green), and Mitotracker (blue). Note formation of numerous membrane ruffles in *PB2/PD1* DKO cells visible by staining with NR12A (red) dye.

**Video S4.** Additional example of SD2 control (left) and *PB2/PD1* DKO (right) GSCs cultured on laminin coated dishes. Note presence of thin long projection from DKO cells stained for Mitotracker (blue) and SPY-tubulin (green). Cell membrane is stained with live cell dye NR12A (red).

**Video S5.** Additional example of SD2 control (left) and *PB2/PD1* DKO (right) GSCs cultured on laminin coated dishes. Note withdrawal of TM-like thin long process from *PB2/PD1* DKO cell, with apparent labeling with Mitotracker (blue) at tip. Cells are also stained with live dyes SPY-tubulin (green) and NR12A (red).

**Video S6.**

SD3 control (top) and *PB2/PD1* DKO GSCs (bottom) migrating through microchannels. Cells were stained with live dyes Memglow (green), SPY-actin (red), and Nucspot (blue). Note presence of cell debris from *PB2/PD1* DKO cells in microchannels.

**Legends for Supplementary Tables**

**Table S1. DEGs of *PLXND1*$^{high}$ vs. *PLXND1*$^{low}$ GBM patients from Rembrandt GBM database.**
Stratification by median expression of *PLXND1*. Cutoff values of adj.$P$<0.01 and $|log_2FC|$>0.58.

**Table S2. DEGs of GBM cells in SD3 *PD1* KO vs. control xenografts, from snRNA-seq data.**
Cutoff values of adj.$P$<$10^{-20}$ and $|log_2FC|$>0.58.

**Table S3. DEGs of SD3 *PB2/PD1* DKO vs. control GSCs, from bulk RNA-seq data.**
Cutoff values of adj.$P$<$10^{-20}$ and $|log_2FC|$>1.

**Table S4. DEGs of SD2 *PB2/PD1* DKO vs. control GSCs, from bulk RNA-seq data.**
Cutoff values of adj.$P$<$10^{-20}$ and $|log_2FC|$>1.

**Table S5. Shared DEGs (*PB2/PD1* DKO vs. control) between SD2 and SD3 GSCs.**

**Table S6. DEGs of TAMs in SD3 *PD1* KO vs. control xenografts, from snRNA-seq data.**
Cutoff values of $P$<0.01 and $|log_2FC|$>0.58.

**Table S7. DEGs in SD3 *PD1* KO vs. control GSCs, from bulk RNA-seq data.**
Cutoff values of adj.$P$<$10^{-20}$ and $|log_2FC|$>0.58.

**Table S8. DEGs in SD3 *PB2* KO vs. control GSCs, from bulk RNA-seq data.**
Cutoff values of adj.$P$<$10^{-20}$ and $|log_2FC|$>0.58.

**Table S9. Shared DEGs of SD3 *PD1* KO vs. control and SD3 *PB2* KO vs. control.**
